# Identifying the genetic basis and molecular mechanisms underlying phenotypic correlation between complex human traits using a gene-based approach

**DOI:** 10.1101/2021.02.09.430368

**Authors:** Jialiang Gu, Chris Fuller, Peter Carbonetto, Xin He, Jiashun Zheng, Hao Li

**Author notes:** send correspondence to and. Both authors contributed equally to this work (Jialiang Gu, Chris Fuller).

## Abstract

Phenotypic correlations between complex human traits have long been observed based on epidemiological studies. However, the genetic basis and underlying mechanisms are largely unknown. Here we developed a gene-based approach to measure genetic overlap between a pair of traits and to delineate the shared genes/pathways, through three steps: 1) translating SNP-phenotype association profile to gene-phenotype association profile by integrating GWAS with eQTL data using a newly developed algorithm called Sherlock-II; 2) measuring the genetic overlap between a pair of traits by a normalized distance and the associated p value between the two gene-phenotype association profiles; 3) delineating genes/pathways involved. Application of this approach to a set of GWAS data covering 59 human traits detected significant overlap between many known and unexpected pairs of traits; a significant fraction of them are not detectable by SNP based genetic similarity measures. Examples include Cancer and Alzheimer’s Disease (AD), Rheumatoid Arthritis and Crohn’s disease, and Longevity and Fasting glucose. Functional analysis revealed specific genes/pathways shared by these pairs. For example, Cancer and AD are co-associated with genes involved in hypoxia response and P53/apoptosis pathways, suggesting specific mechanisms underlying the inverse correlation between them. Our approach can detect yet unknown relationships between complex traits and generate mechanistic hypotheses and has the potential to improve diagnosis and treatment by transferring knowledge from one disease to another.

## Introduction

The human body is a complex system with multiple interacting components, and as such many traits are inter-related. Phenotypic correlations between complex traits have long been observed based on epidemiological studies. Depending on the level of our understanding of the underlying mechanism, some of the relations may seem obvious, e.g., Fasting Insulin vs. Fasting Glucose, while others may seem totally unexpected, such as the inverse correlation between Cancer and Alzheimer’s disease[1, 2]. It has been argued that many of the phenotypic correlations have genetic underpinning, based on the inheritance patterns of the traits involved. However, little is known due to the polygenic nature of these traits. Understanding phenotypical connections at the genetic level can potentially lead to new mechanistic insight and allow the transfer of diagnosis and treatment from one well-studied disease to another. Such information can also be used to improve polygenic risk prediction[3, 4].

The recent application of GWAS to a wide range of complex human traits has provided an opportunity to systematically identify genetic similarity between traits and delineate the underlying mechanism. Several comparative GWAS analysis tools have been developed to detect genetic similarity between phenotypes based on co-associations at the SNP level, and applications of these tools to GWASs have lent genetic support to some of the earlier epidemiological observations[5–11]. However, the genetic similarity between different traits can be obscured at the SNP level. Phenotypes sharing a set of intermediate molecular traits such as gene expression may not necessarily associate with the same SNPs, since multiple SNPs can converge on the same gene, and only a small subset of these SNPs may be identified in a particular GWAS. Furthermore, it is challenging to infer common molecular mechanisms based on the shared SNPs, as they often fall into non-coding regions with no apparent functional implication. Therefore, it is logical to consider genetic similarity at the gene level that integrates information from multiple convergent SNPs.

Since the majority of GWAS identified SNPs are in the non-coding regions of the genome, where they often overlap with regulatory elements such as enhancers, promoters, and transcription factor binding sites, which influence the expression levels of nearby or distant genes [12–14], it is natural to consider integrating information from multiple SNPs that influence the expression of the same gene. Such information can be derived from the systematic eQTL analysis of multiple human tissues.

Here we introduce a gene-based approach to quantify the extent to which two complex traits share similarity in the genes they associate with, defined as genetic overlap between the traits. For a given GWAS, we first integrate the information of all SNPs that may regulate the expression of a gene (using eQTL data) to derive a p value of association between the gene and the phenotype, and then treat the gene-phenotype p values for all the genes as a genetic profile of the phenotype. We then analyze the overlap between the genetic profiles of two different phenotypes and calculate a genetic overlap score and assess the statistical significance. Once a significant overlap between two phenotypes is detected, we identify groups of functionally related genes (such as those defined by gene ontology (GO) terms or KEGG pathways) that contribute to the overlap using the Partial Pearson Correlation Analysis (PPCA) that we previously developed [15]. This generates hypotheses regarding common physiological processes underlying different and sometimes seemingly unrelated phenotypes.

Using this approach, we analyzed a dataset of ∼400 GWASs, using a combined eQTL dataset, including GTEx[16]. We measured genetic overlap between 59 phenotypes and identified many known and unexpected relationships. Many of the discovered relationships are supported by epidemiological observations. Examples include Alzheimer’s Disease (AD) and cancer, Rheumatoid Arthritis and Crohn’s disease, and Longevity and Fasting Glucose. Our approach can detect genetic overlap between seemingly unrelated phenotypes that are not detectable by SNP-based similarity measures[8], and importantly, for each of the detected pairs, our method suggests a number of shared genes/pathways that point to potential mechanistic connection between the two phenotypes.

## Results

### Method Overview

We formulate the problem as given a GWAS (query GWAS), search for a database of GWASs for significant matches (hit GWAS), based on their gene-phenotype association profiles (Figure 1). This is similar in spirit to BLAST searches for sequence similarity. Starting with GWAS data, we assign a p-value of association for each gene with a given phenotype, so that each phenotype is represented by a gene association vector of dimension N, while N is the number of genes in the genome. This step translates the SNP-phenotype association profile from the original GWAS into a gene-phenotype association profile. For this purpose, we developed a new computational algorithm called Sherlock-II, to integrate GWAS with eQTL data using the collective information of all SNPs. We then score the “genetic overlap” between two phenotypes based on the similarity of the two gene-phenotype association profiles, using a normalized distance between the two, we called it “genetic overlap score” Sg. We assess the statistical significance of the measured genetic overlap score between a query GWAS and a hit GWAS using a background distribution generated by an ensemble of random GWASs with the same power as the hit GWAS (Figure 1 and Methods). This second normalization step normalized out the dependence of Sg on the power of GWASs being compared, resulting in a z score Z_S_, which can be assigned a p value. Query-GWAS/hit-GWAS pairs with significant p value indicates that the two GWASs have significant genetic overlap at the gene level. We found that the signal is often distributed to many genes, instead of a small number of top genes associated with both phenotypes.

**Figure 1:**
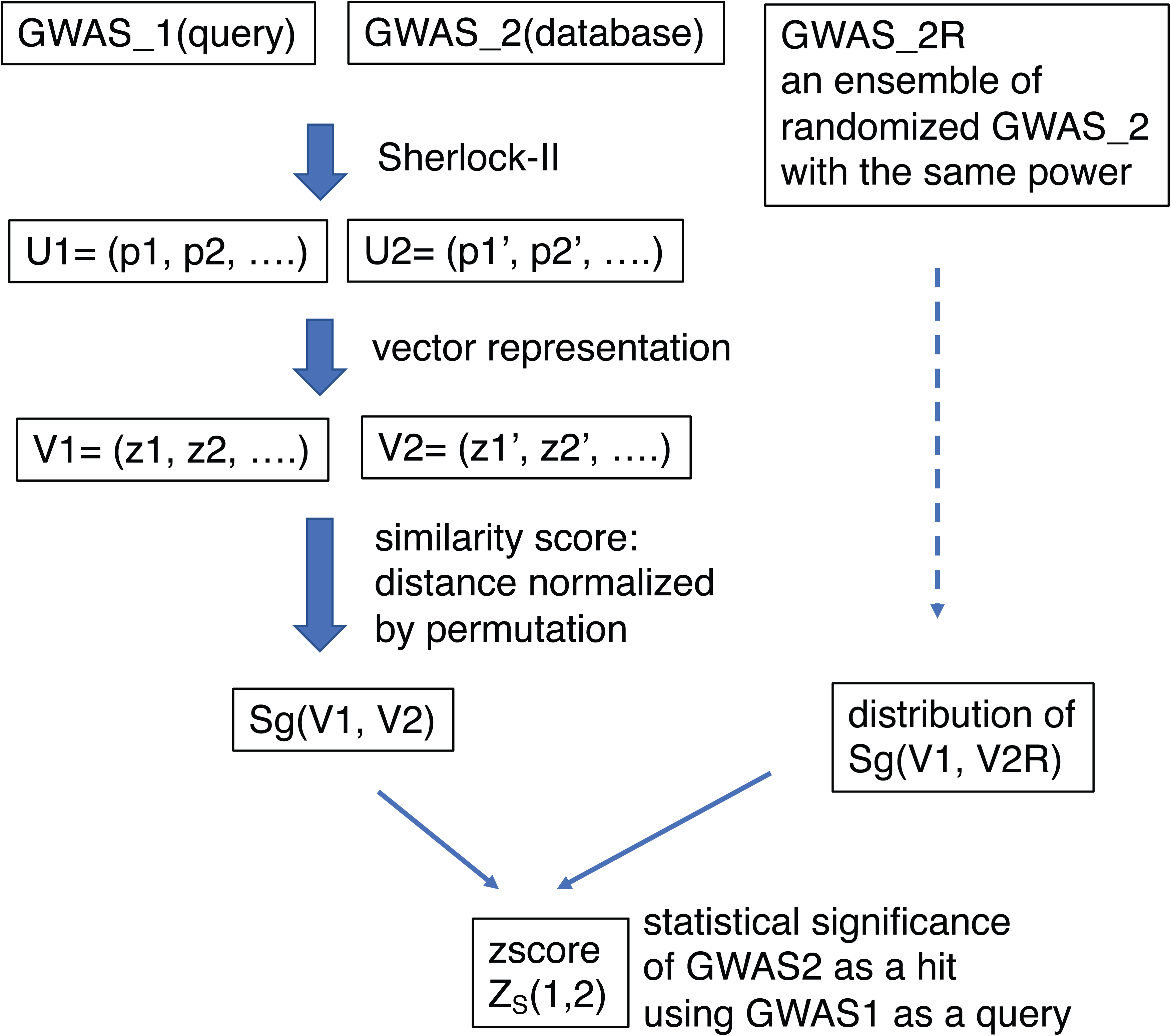
An overview of the gene based approach for detecting genetic overlap between complex traits. Starting from a pair of GWASs (GWAS_1 and GWAS_2) representing two complex traits, this approach first transforms the GWAS data into gene-phenotype association profiles (vectors U1, U2), by assigning a p value for each gene using Sherlock-II (through integration of GWAS and eQTL data). The gene-phenotype p values are then translated into unsigned z scores (vectors V1, V2), and a gene based genetic overlap score Sg(V1, V2) is calculated as a permutation normalized distance between V1 and V2. To assess the statistical significance of the score Sg, an ensemble of random GWASs with the same power as GWAS_2 is generated (GWAS_2R), and the score Sg(V1, V2R) between GWAS_1 and GWAS_2R is calculated in parallel (see Methods). The distribution of Sg(V1, V2R) is then used to normalized the score Sg(V1, V2), leading to a final genetic overlap z score Z_S_(1,2), from which a p value can be calculated.

### Translate SNP-phenotype association profile to gene-phenotype association profile using Sherlock-II algorithm

The first step in the gene-based approach is to assign a p-value of association for each gene to a given phenotype, thus translating the SNP association profile of the original GWAS to a gene association profile. To have better power, such transformation should preserve as much information from the GWAS as possible (not just the SNPs passing the genome-wide significance threshold) and utilize the convergent effect of multiple SNPs on the same gene. Thus, we need a high throughput tool to combine GWAS data with eQTL data that can capture the collective information from multiple SNPs related to a gene, both in cis and trans. In addition, due to the scale of computation, this tool should be statistically robust and computationally efficient.

Multiple computational tools have been developed to associate complex traits with genes [17–24]. However, most of the existing tools infer the association between gene and trait by colocalizing the GWAS and eQTL signal in a single locus, starting from a strong GWAS or eQTL peak. Such an approach leads to predictions with high specificity but may miss genes supported by the alignment of multiple GWAS and eQTL peaks that do not pass the genome-wide significance threshold. Previously, He et al. developed the Sherlock algorithm that use the collective information of all the SNPs to infer gene-phenotype association, including both cis and trans eSNPs, thus performing global alignment between the GWAS and the eQTL profiles[20]. However, the Bayesian formulation used makes it difficult to compute p-values and is sensitive to inflation in the input data. Here we developed a new computational algorithm called Sherlock-II, a second generation of Sherlock that overcomes the limitations of Sherlock, to integrate GWAS with eQTL data.

Similar to Sherlock, Sherlock-II detects gene-phenotype associations by comparing the GWAS SNPs with the eQTL SNPs of a gene, with the assumption that if the gene is causal to the phenotype, SNPs that influence the target gene’s expression should also influence the phenotype; thus the eQTL profile of the gene should have significant overlap with the GWAS profile of the phenotype. Although conceptually similar, Sherlock-II uses a totally different statistical approach to calculate the statistical significance of the profile alignment, leading to a number of significant improvements. While Sherlock calculates a Bayesian factor for the observed eQTL and GWAS profiles and uses randomization of GWAS data to estimate p-value, Sherlock-II uses a statistic that is a sum of the log(p-value) of the GWAS peaks aligned to the eQTL peaks, with background distribution of the statistic directly calculated by the convolution of the distribution of the log(p-value) for all the independent GWAS peaks (see Methods). This new approach gives Sherlock-II several advantages: 1) it is more robust against inflation. By deriving the background distribution empirically from the list of all p-values from the GWAS SNPs aligned to tagged eSNPs (in independent LD blocks), it automatically accounts for inflation that may exist in the original GWAS data; 2) it gives more accurate and efficient p-value calculation since the background distribution can be calculated through convolution, instead of GWAS randomization. These combined features of Sherlock-II made it possible to perform a global analysis of GWASs vs. eQTL data efficiently.

To further demonstrate the improvement of Sherlock-II over Sherlock for integrating GWAS with eQTL data, we generated synthetic data with known ground truth, by picking a number of causal genes, and for each gene assign a variable number of eSNPs (passing a threshold) as causal SNPs (mediated through the gene), sample the effect size, and simulate the GWAS data using the simGWAS[25] algorithm. We then apply Sherlock-II and Sherlock to the simulated GWAS data and the real eQTL data to infer the causal genes. This comparative analysis demonstrated that Sherlock-II performs better than Sherlock, due to reduced false positive rate, as it uses a better background model. Sherlock-II is also 30 times faster if the desired lowest p value is 10^(-6), which requires 10^6 simulations for Sherlock (Figure S1).

Using Sherlock-II, we have analyzed a set of the published GWAS data, using the eQTL data we curated. The analyses include 445 GWAS datasets covering 88 phenotypes, 8 individual eQTL covering 6 unique tissues[26–33] and GTEx (version 7) data[34]. To include the possibility that the expression of a gene can influence the phenotype through multiple tissues, we combined eQTL data across different tissues: when a SNP appears in multiple eQTL datasets, we assign the most significant p-value to it to capture the best eQTL signal across tissues. This merged version includes the eQTL profile of 21,892 genes. For each GWAS dataset, Sherlock-II calculates a p-value for every gene measuring the statistical significance of the overlap between the merged eQTL profile of the gene and the GWAS profile, thus translating the original SNP-phenotype association profile to a gene-phenotype association profile. Overall, we have identified 763 gene-phenotype associations with FDR <0.05, covering 317 genes and 52 phenotypes (Dataset S1). Many of these findings have literature support and are biologically meaningful.

In many cases, although only weak SNP associations are present in the GWAS data, Sherlock-II detects strong gene-phenotype associations as many GWAS SNPs are considered in aggregate during the transformation. These strong associations are often supported by multiple aligned SNPs with weak to moderate GWAS p values, and collectively they give a strong signal. Examples include the autism GWAS, of which the original signal is weak (no SNP passing genome-wide significance threshold)[35]. Sherlock-II identified 10 genes with FDR<0.1; four of them (ENPP4, MAPT, LEPR, MTHFR) have literature support[36–39] (Figure S2).

We have also applied Sherlock-II to metabolite-QTL data[40] and discovered 22 metabolite-phenotype associations at FDR<0.05, covering 20 metabolites and 9 phenotypes (Dataset S1). Sherlock-II identified several metabolites with a strong association with neuro-degenerative/neuro-psychiatric diseases, even though the signal in some of the original GWAS data is very weak (Figure S3).

### Measure genetic overlap between two phenotypes using the global gene association profiles and identify shared genes/pathways

The next step in the gene-based approach is to score the genetic overlap between two phenotypes based on their gene association profiles. Starting with the p-value of association between a gene *i* and a trait *t*: *P_i_*(*t*), we pick a vector representation for the association profile of the trait *t*, *V*(*t*) = [*Z_1_*(*t*), *Z_2_*(*t*), …, *Z_N_*(*t*)] where *Z_i_*(*t*) is the z score corresponding to the p-value *P_i_*(*t*), based on a two tailed test using standard normal distribution (see Methods), and *N* is the total number of genes in the genome. We use an unsigned z score because we do not consider the direction of the gene’s effect on the phenotype (see Discussion). We then measure the Euclidian distance between the two phenotypes E(*V*(*t*_1_),*V*(*t*_2_)). We expect that phenotypes with shared genes will have certain components of the vectors with similar values, and thus will give rise to a smaller distance. We define a genetic overlap score between the two traits as 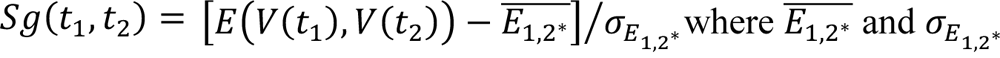 are the mean and standard deviation of the Euclidean distance generated by randomly permuting the components of the second vector. For two phenotypes with genetic overlap, we expect that the distance will be smaller than the mean, thus Sg will be negative. Trait pairs with more overlap will have a more negative Sg. This distance normalization is important as it takes out the contribution to the distance due to the features of each individual vectors, since we are interested in the overlap between the two vectors, not the vectors themselves. If the two vectors are un-correlated, the score defined above should have a standard normal distribution and therefore can be assigned a statistical significance. However, the components of the two vectors are correlated even if one of the GWASs is random, due to the structure of the eQTL data. E.g., pleiotropic eSNPs will lead to the correlated fluctuation of the p values of their target genes, and such correlation is stronger for higher powered GWAS (Figure S4).

To calculate the statistical significance of the observed genetic overlap score Sg, we consider comparing a query GWAS to a database of GWASs to identify which one has a significant overlap with the query - called it a hit GWAS. We define P(S) as the probability by chance that the query GWAS and a randomized GWAS with the same power as the hit GWAS will have a score less than or equal to Sg. By using an ensemble of random GWASs with the same power as the hit GWAS, we generate a null distribution of the overlap scores between the given query GWAS and the random GWASs. We then use the null distribution to perform the second normalization, resulting in a score Z_S_ that follows z distribution, and thus can be used to calculate p value (Figure 1 and Methods). Using random GWASs with the same power as the hit GWAS to normalize Sg is necessary to control for the dependence of Sg on the power of hit GWAS (Figure S4). Note that although the genetic overlap score is symmetric with respect to the two GWASs, the p value defined above is not symmetric, i.e., switching the query-GWAS and hit-GWAS can lead to a different p values. However, in practice, if we find a pair with significant overlap, it is generally significant in both directions.

We verified that the p values and the ranking of the hit GWASs is not sensitive to the actual vector representation and the distance metric we use. For example, using -log_10_(*P_i_*(*t*)) as the vector component or using Manhattan distance as a distance metric gives very similar results (see Figure S5, 6).

One major advantage of gene-based approach over SNP-based methods is that once the genetic overlap is detected, the information on genes/pathways that support the overlap can be readily delineated. A straightforward approach is to select the top genes for each trait with a p-value cutoff and analyze the overlap. This approach does yield interesting results for some trait pairs. However, for the majority of trait pairs that we detected, the top associated genes for the two traits do not overlap. This suggests that the genetic overlap is driven by a more diffused signal distributed to a broad set of genes. To identify such set of genes, we employ an approach we previously developed to identify functionally related genes supporting the similarity between two gene expression profiles[15]. This approach assesses the contribution of a subset of genes (e.g., defined by a GO category or a KEGG pathway) to the global Pearson correlation and assigns a z-score to the subset (see Methods); we called it Partial-Pearson-Correlation Analysis (PPCA). This allows the ranking of the significance of different subsets of genes based on their z-score (PPCA-zscore). This approach enables us to identify sets of GO terms and KEGG pathways that significantly contribute to the genetic overlap between a pair of traits and generate hypotheses regarding the mechanisms that connect the two traits.

### Application of gene-based approach to 59 Human traits revealed many known and unexpected relationships and potential mechanisms

Using the approach described above, we analyzed the relationships between 59 Human traits covered by the 69 previously published GWAS data. This is a filtered version of the 445 GWAS datasets analyzed by Sherlock-II, with redundant data and many sub-phenotype groupings in the same GWAS removed. For a few traits, we kept multiple GWAS studies published separately as positive controls.

Starting from the gene-association profiles generated by Sherlock-II, we calculated the genetic overlap scores and p values using each of the 69 GWASs as the query against other 68 GWASs. We detected significant overlap for 1346 pairs with Zs below -4 (p value =3.17e-5, expect 0.15 positives from all the pairwise comparisons). Here we present the top hits for a selected subset of 12 traits in Figure 2, including some highly prevalent diseases such as cancer, AD, and Diabetes. The full result of the 69x68 comparisons is provided online (Dataset S2).

**Figure 2.**
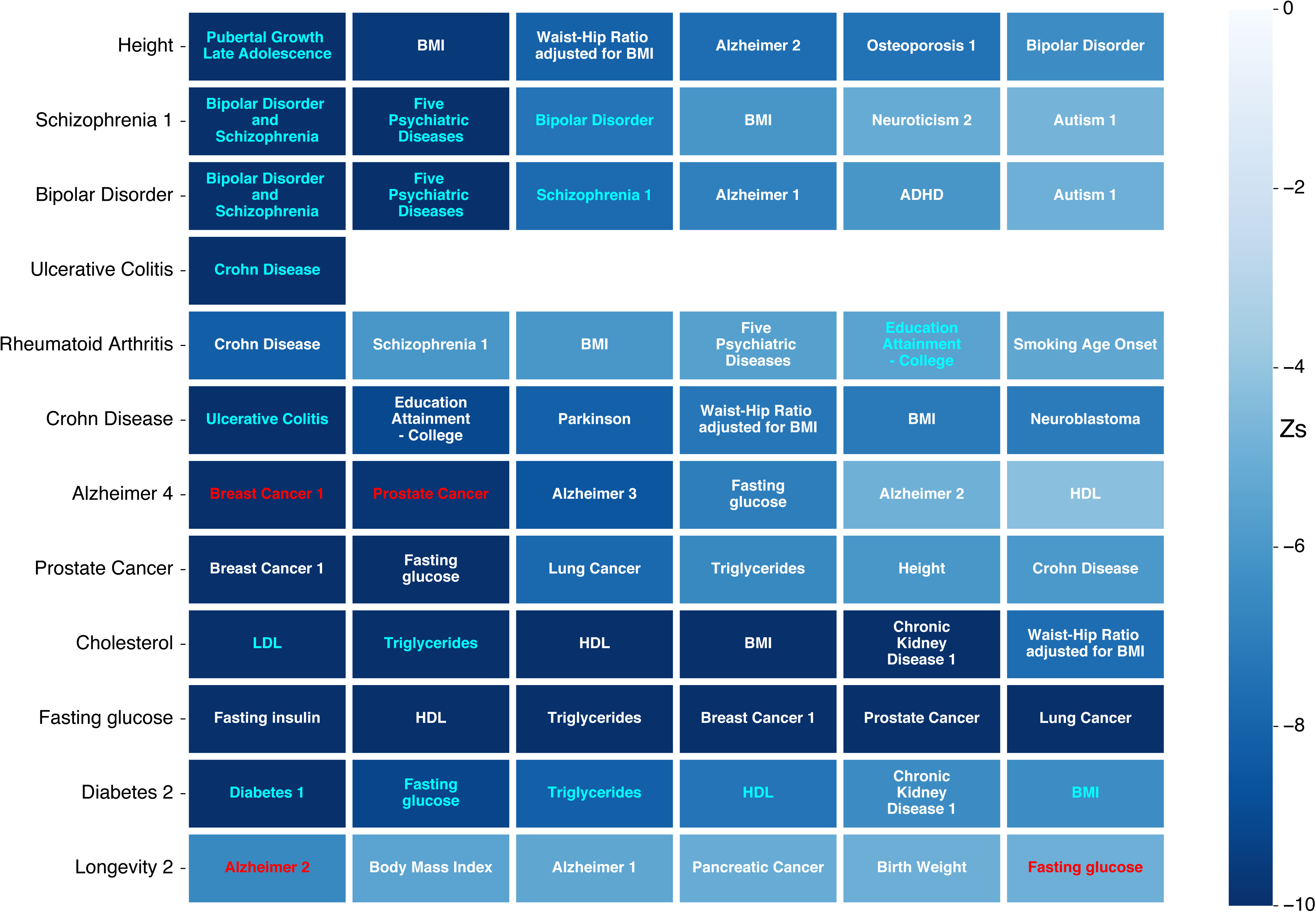
Genetic overlap between pairs of traits detected by the gene-based approach. Shown are top hits to 12 selected traits (query GWAS, first column). Each row listed the top 6 hit GWASs that have Z_S_ < -4 with the query GWAS (pvalue < 3.17e-5), ranked by their p-values (uncorrected for multiple test) as indicated by the color scale. Pairs detected in a previous analysis using LDSC with the same p value cutoff (p=3.17e-5, uncorrected for multiple test) are colored in cyan, and the examples of unexpected relationships discussed in the text are colored in red. Different numbers associated with the same trait name correspond to different GWAS data (pubmed ID) for the trait: Alzheimer 1(PMID 17998437), Alzheimer 2(PMID 21460841), Alzheimer 3(PMID 24162737), Alzheimer 4(PMID 25188341), Breast Cancer 1(PMID 17903305), Breast Cancer 2(PMID 29059683), Chronic Kidney Disease 1(PMID 20383146), Diabetes 1(PMID 18372903), Diabetes 2(PMID 22885922), Neuroticism 2(PMID 27089181), Osteoporosis 1(PMID 22805710), Schizophrenia 1(PMID 25056061), (see Dataset S3 for the full mapping from a trait name to the GWAS data).

Many of the detected pairs are known or biologically meaningful based on our understanding of the relationship between the two traits. Besides obviously related pairs of traits, such as Diabetes vs. Fasting Glucose or Height vs. Body Mass Index (BMI), our method detected genetic overlap between traits known to share similar etiology. For example, Crohn’s disease vs. Ulcerative Colitis, Schizophrenia vs. Bipolar Disorder, and Fasting Glucose vs. Fasting Insulin. Many of these relations were also detected previously through comparative GWAS analysis using a well-established SNP based similarity measure-LD score regression (LDSC) [8] (Figure 2, entries highlighted in cyan color).

Our method also detected a number of pairs expected to share similar etiology but not detected by LDSC. One such example is the relation between RA and CD (Z_S_= -8.16, p-value=1.7e-16 when RA is the query, and Z_S_ = -6.45, pvalue=5.6e-11 When CD is the query) known to share inflammatory/autoimmune origin (LDSC gives a regression coefficient r = -0.029 and p-value= 0.7255). Using PPCA of GO and KEGG terms (see Methods), we identified a number of gene groups shared by the two traits, with a strong theme in inflammation and immune responses, including MHC class II protein complexes and pyroptosis, which is a highly inflammatory form of programmed cell death, supporting the idea that cell death caused by dysregulated inflammatory response is a common etiology for the two diseases (Figure 3) [41, 42].

**Figure 3.**
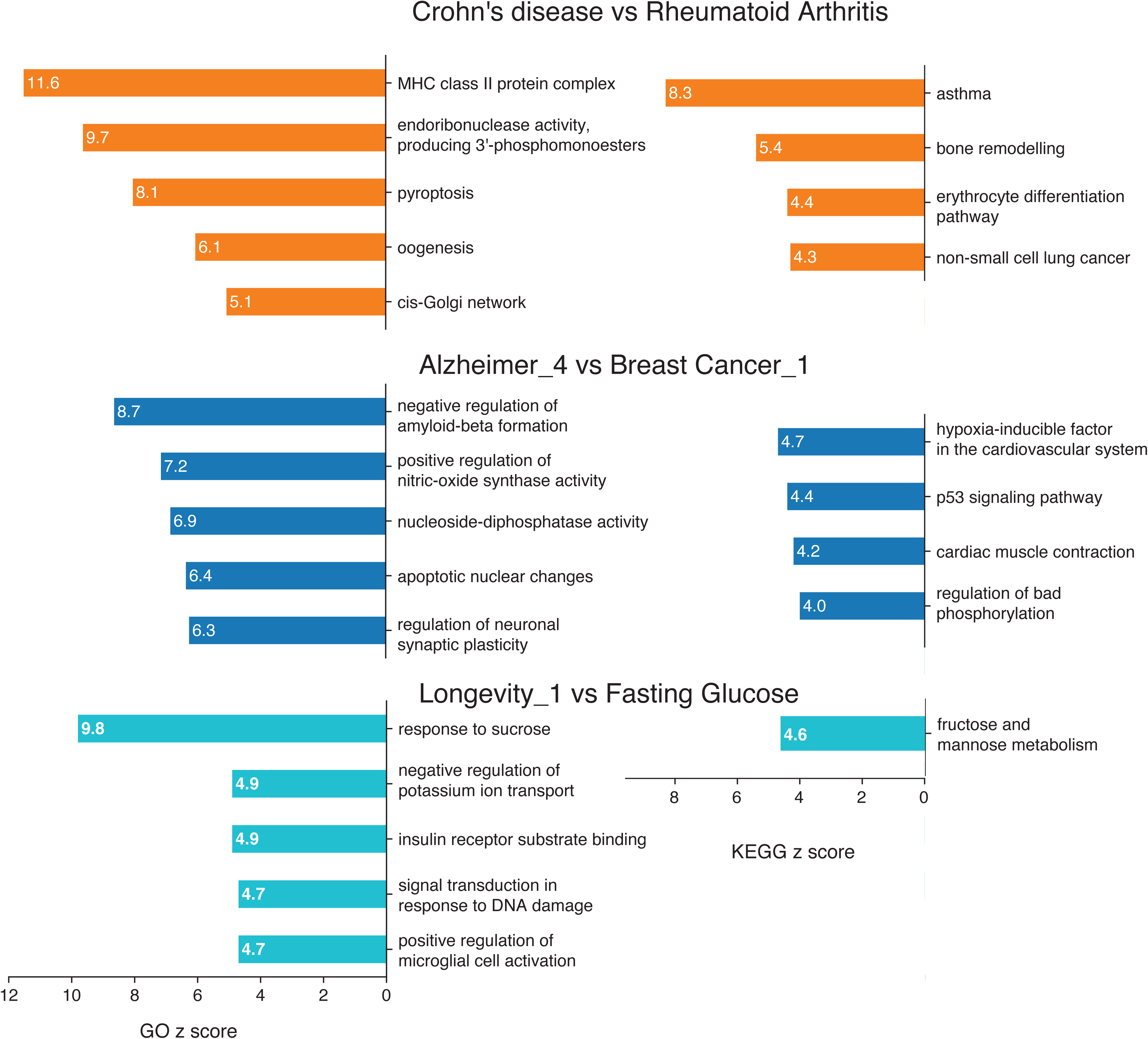
Partial Pearson Correlation Analysis (PPCA) revealed GO terms and KEGG pathways supporting the genetic overlap between traits. Shown are GO terms with PPCA-zscore >5 and KEGG pathways with PPCA-zscore >4, up to top 5, for Crohn’s Disease vs. Rheumatoid Arthritis, AD vs. Breast Cancer, and Longevity vs. Fasting glucose pairs. Length of the bar and the numbers in the bar indicate PPCA-zscore.

Importantly, our gene-based approach identified a number of unexpected connections between seemingly unrelated traits. Many of these relationships have support from previous epidemiological studies, but the genetic basis and mechanism for the connections are largely unknown. We discuss two examples in more details below.

### Genetic overlap between AD and Cancer and the supporting genes/pathways

Epidemiological studies found that risk for cancer and AD is inversely correlated. E.g., in a prospective longitudinal study, it was found that the risk of developing cancer is less among participants with AD vs. nondemented participants and that the risk of developing AD is less for participants with a history of cancer[2]. A recent study of the association between cancer and longitudinal progression of dementia also showed inverse correlation[1]. It has been speculated that such inverse correlation may involve genes regulate cell growth vs. maintenance and cell survival vs. cell death[43]. However, the genetic basis for the correlation and specific genes/pathways that support the correlation have not been analyzed systematically.

We detected a strong overlap between AD and breast and prostate cancer, with p values of =5.8e-35 and 6.2e-23 (Z_S_ = -12.28, -9.79) respectively (Figure 2).

We applied PPCA to the AD-Breast Cancer pair, and identified 8 GO terms and 4 KEGG pathway with PPC-zscore > 4 (Figure 3). The top 5 GO terms (representative genes) are “negative regulation of amyloid-beta formation” (APOE, BIN1), “positive regulation of nitric-oxide synthase activity” (APOE, AKT1, HIF1A), “nucleoside-diphosphatase activity” (ENTPD6, NUDT5), “apoptotic nuclear changes” (SHARPIN), and “regulation of neuronal synaptic plasticity” (APOE, SYT4). Consistent with the GO terms, two most significant KEGG pathways were “hypoxia-inducible factor in the cardiovascular system” (P4HB, HIF1A) and “p53 signaling pathway” (SHISA5, CCND3). Several of the representative genes were previously found to be associated with either AD or cancer, or both. E.g., APOE is the most famous gene for AD, and is also associated with cancer[44]. AKT1 has been reported to govern breast cancer progression in vivo[45] and reactive oxygen species-mediated loss of synaptic Akt1 signaling leads to deficient protein translation early in AD[46].

Two functional themes were found by the PPCA. One is hypoxia response, implicated by both GO (“positive regulation of nitric-oxide synthase activity”) and KEGG (hypoxia-inducible factor in the cardiovascular system). Nitric oxide (NO) is a major signaling and effector molecule mediating the body’s response to hypoxia[47]. It is known that tumors grow in an environment that lacks oxygen, and the ability to adapt to a hypoxic environment is a key to cancer survival and proliferation. Hypoxia is also linked to neural inflammation, and cerebral hypoxia has been linked to beta-amyloid formation in AD patients. A better hypoxia response may lead to the suppression of AD pathology[48]. Thus, hypoxia response may serve as one mechanism underlying the inverse correlation between cancer and AD. The second functional theme is p53 signaling and apoptosis, also implicated by both GO and KEGG. P53 is a master regulator of cell fate, including cell cycle, senescence, and apoptosis. It is a cancer suppressor that is most frequently mutated in various cancer cells. Genetic variations that strengthen P53 signaling will better suppress cancer. However, in the context of AD, it may also increase the neuronal death that is a major feature of AD pathology. Thus, P53 pathway/apoptosis may be another mechanism supporting the inverse correlation between cancer and AD.

### Longevity and Fasting Glucose

Comparative analyses of GWASs of human longevity and other traits have identified overlap between lifespan and a number of age related diseases, including AD, coronary artery diseases, and cancer [49]. In an earlier GWAS of centenarians, a SNP in the ApoE locus stands out as having the most significant association, a result that has been replicated in several subsequent longevity GWASs, linking extreme longevity to AD.

We found that a meta GWAS of long lived individual (> = 90 years) with European descent (cases n=5406, controls (<65 years) n=15 112)[50] has significant similarity to AD and pancreatic cancer with p values of 2.0e-11 and 1.7e-7 (zscore of -6.6 and -5.1) respectively (Figure 2).

Interestingly, this longevity GWAS also shares similarity with fasting glucose (p value =4.8e-7; z = -4.9). Fasting glucose is strongly correlated with fasting insulin, and insulin and insulin like growth factor (IGF) signaling has been implicated in regulating lifespan across multiple model organisms [51]. However, GWASs of longevity have not been able to identify insulin/IGF signaling pathway genes, although targeted sequencing of IGF1 and IGF1 receptor genes did reveal enrichment of rare variants in the centenarian cohort[52]. This suggests that mutations in this pathway extend lifespan but may have deleterious effect on development (i.e., antagonistic pleiotropy); thus, only those with small effect size can become common variants included in GWAS. Our method identified the relation between fasting glucose and longevity by integrating information from many SNPs with small effect size.

To explore potential causal relationship between longevity and fasting glucose, we applied PPCA to the pair and identified 5 GO terms and one KEGG pathway with PPCA-zcore >4 (Figure 3). The top three GO terms (representative genes) are “response to sucrose” (KHK, FADS1), “negative regulation of potassium ion transport” (HCRT), and “insulin receptor substrate binding” (WDR6,GRB2). The significant KEGG pathway is “fructose and mannose metabolism” (KHK, PFKFB3). Thus, both GO and KEGG point to sugar metabolism and insulin signaling. Since these shared genes/pathways relate more directly to fasting glucose, a more likely causal model is fasting glucose -> longevity, instead of longevity -> fasting glucose, consistent with the genetic studies in model organisms.

### Clustering of phenotypes based on their genetic overlap scores

As described above, using a query GWAS to search for a database of GWASs allows us to identify related traits, much like a BLAST search for sequence similarity. A different perspective on this data is to cluster phenotypes together based on their genetic overlap by mapping the genetic overlap scores (Z_S_ (1,2) and Z_S_ (2,1)) to a distance that is symmetric (see Methods). The result of this global clustering analysis is a tree in which phenotypes with high overlap scores are placed next to each other in a subtree (Figure 4). The advantage of this approach is that the tree will capture the relationships between the multiple phenotypes in the clusters, although significant pairwise overlap may not be readily visible due to the hierarchical nature of the clustering.

**Figure 4.**
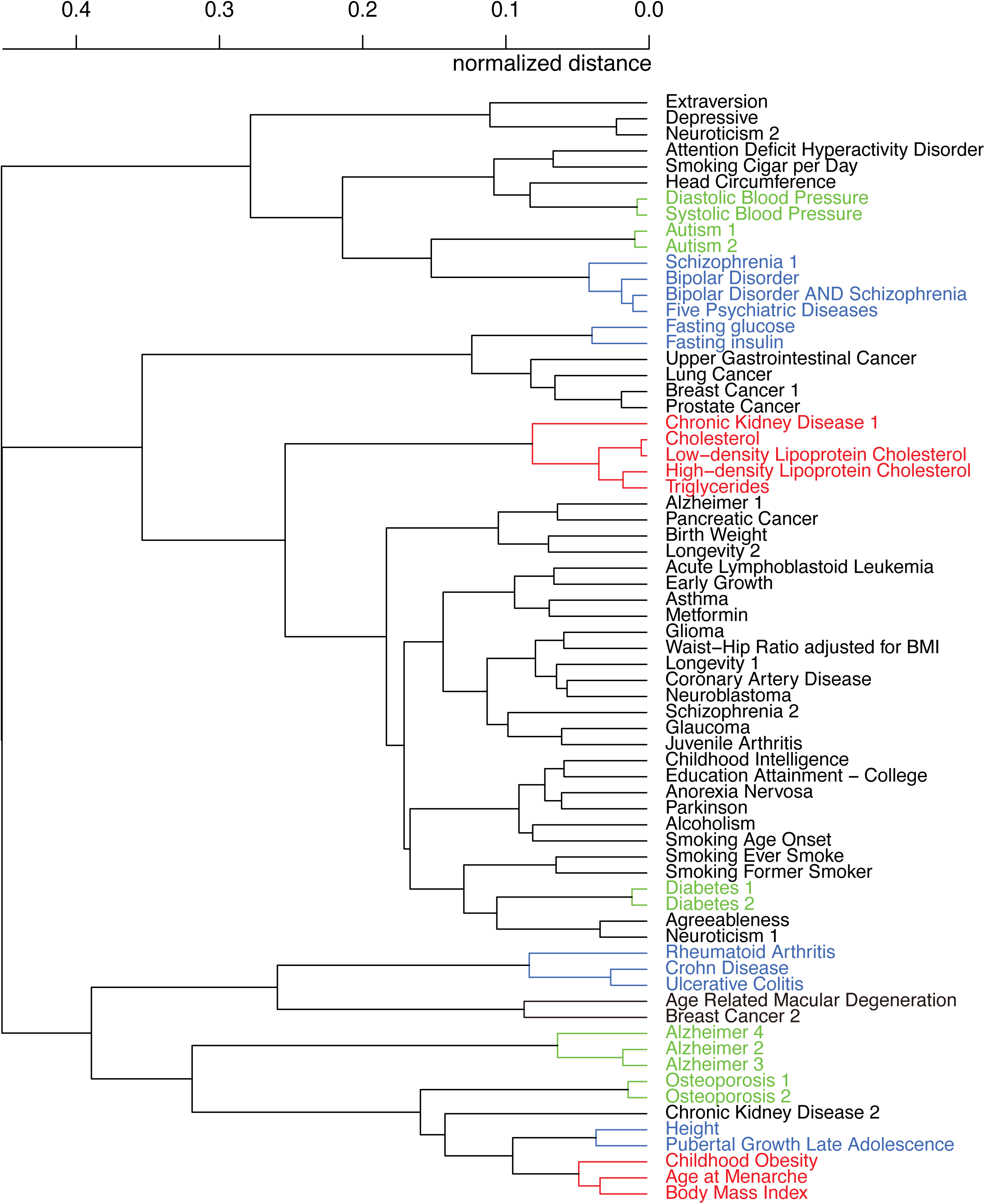
Hierarchical clustering of phenotypes based on their pairwise genetic overlap score. Highlighted branches discussed in the text: 1) GWASs of the same phenotypes (green); 2) phenotypes that are known to share genetic etiology (blue); and 3) phenotypes with no obvious relations but co-occurrences were observed based on previous epidemiological studies (red). The colored subtrees were picked by cutting the tree at a distance of 0.2 (average Z_S_ <-4).

We found that similar/related phenotypes were clustered together as expected. These sub-clusters are formed by 1) GWASs of the same phenotype (e.g., osteoporosis, AD, diabetes, and autism); 2) phenotypes that are closely related and were studied in the same GWASs (e.g., LDL cholesterol, HDL cholesterol, total cholesterol, and Triglyceride; diastolic and systolic blood pressure); 3) phenotypes that are known to share genetic etiology in previous studies (e.g., bipolar disorder (BP), schizophrenia (SZ), the combination of the BP and SZ, and five psychiatric disorders including BP and SZ [53], and 4) phenotypes with strong evidence supporting their shared mechanisms (e.g., Rheumatoid arthritis, Crohn’s disease, and Ulcerative Colitis, due to their common inflammatory/autoimmune nature).

The clustering analysis also revealed a number of sub-trees in which phenotypes with no obvious relations are linked together (Figure 4). Here we discuss two examples: 1) a subtree that contains age at menarche, childhood obesity, body mass index; 2) a subtree with 5 phenotypes: cholesterol, HDL, LDL, triglycerides, and chronic kidney disease (CKD). Upon literature search, we found that the groupings were supported by previous epidemiologic observations: 1) a previous study found that girls experiencing early menarche are significantly overweight/obese [54]; 2) it was found that people with high total cholesterol or low HDL were more likely to have reduced glomerular filtration rate, which is the canonical way to assess kidney function[55], and it is not surprising to see total cholesterol, HDL, LDL and triglycerides are clustered together, as they are generally correlated.

We next analyzed the shared genes/pathways between the phenotypes in these subtrees. For the age-at-menarche/body-mass-index/childhood-obesity subtree, we found overlapping genes in each of the three pairwise comparisons. Multiple genes are associated with age-at-menarche and body-mass-index with a theme on cell growth and proliferation. For example, SMAD3 is significantly associated with both age-at-menarche and body-mass-index and is a pivotal intracellular nuclear effector of the TGF-beta (transforming growth factor beta) signaling which plays a critical role in regulating cell grow, differentiation and development[56]. MAP2K5 (Mitogen-Activated Protein Kinase Kinase 5) is also significantly associated with both the phenotypes and the signaling cascade mediated by this kinase is involved in growth factor stimulated cell growth and muscle differentiation [57]. Genes simultaneously associated with both childhood-obesity and body-mass-index include TFAP2B and CENPO. TFAP2B is a transcription factor of AP-2 family that stimulates cell proliferation and suppresses terminal differentiation of specific cell types during embryonic development [58]. CENPO encodes a component of the interphase centromere complex which is required for bipolar spindle assembly, chromosome segregation, and checkpoint signaling during mitosis[59]. Thus, these traits share a set of genes with a common theme on cell growth/division and development.

The connection between cholesterol and CKD was observed in epidemiological studies and often interpreted as that high cholesterol is hazardous to kidney function. We performed PPCA to identify shared functional gene groups that support the connection. Interestingly, between CKD and any of the other four phenotypes, we did not see any obvious functional groups that directly link to cholesterol catabolism/homeostasis. Instead, one of the strongest terms is glomerular basement membrane development (Figure S7), which is related to kidney function. Two representative genes in this group are MPV17 and LAMB2. MPV17 encodes a mitochondrial inner membrane protein implicated in the metabolism of reactive oxygen species[60]. Mutation of this gene cause Focal Segmental Glomerulosclerosis[61] in mice, a disease in which scar tissue develops on the parts of the kidneys that filter waste from the blood. LAMB2 is a member of the Laminins family, a family of extracellular matrix glycoproteins that are the major non-collagenous constituent of basement membranes. Mutations in LAMB2 cause Congenital Nephrotic Syndrome in human[62], which is a kidney condition that begins in infancy and typically leads to irreversible kidney failure by early childhood. These observations favor a causality relation (MPV17, LAMB2, ..) -> kidney function -> cholesterol levels, instead of cholesterol -> kidney function as it is generally interpreted. For comparison, we analyze genes shared within the four related phenotypes cholesterol/HDL/LDL/triglyceride, and found that the strongest terms are generally related to cholesterol catabolism/homeostasis, such as “negative regulation of lipoprotein lipase activity”, “phospholipid homeostasis”. Thus, gene based approach can help infer causality for observed phenotypic correlation, in the spirit of Mendelian randomization, if there are clear functional annotations of the genes.

## Discussion

We have developed a computational framework to analyze the genetic overlap between complex traits and delineate the potential mechanisms underlying the overlap by integrating GWAS of multiple phenotypes with eQTL data. We demonstrated the approach by a global analysis of published GWASs covering 59 complex human phenotypes. Our analysis revealed expected as well as unexpected relationships between phenotypes and suggested specific mechanisms that connect these phenotypes.

Compared with genetic similarity analysis methods using GWAS data at the SNP level, our gene-based genetic overlap analysis has several advantages. First, comparing the genetic profiles at the gene level can yield a stronger signal, as multiple SNPs can converge on the same gene. Even if the GWAS peaks of two different phenotypes do not align by proximity, they can be “aligned” through a gene, i.e., their effects on the phenotype can be mediated through the same gene. Thus, our method can be more sensitive in detecting shared genetic basis between traits. Our method identified many trait pairs with a strong overlap that were not detected by SNP based similarity measures such as LDSC regression[8] (Figure 2). Examples include AD vs. breast cancer, which was observed in previous epidemiological studies, and Crohn’s disease vs. Rheumatoid Arthritis, which are known to share inflammation/immune origin. Secondly, genetic overlap analysis at the gene level readily suggests potential mechanisms connecting different phenotypes. We used the PPCA method to identify groups of functionally related genes co-associated with a pair of traits; such analysis led to new insight into the mechanistic connections between the two phenotypes.

We note that our method may also miss some relations detected by LDSC. This can happen, e.g., if the two phenotypes share some GWAS SNPs that do not align to eSNPs, either because the SNPs do not influence the phenotype by influencing mRNA expression, or the eQTL data (which is often noisy and incomplete) failed to capture the eSNPs. In a systematic comparison, we found that among the 1378 pairs (53 unique phenotypes) analyzed by both LDSC and our method, 1118 pairs produce valid LDSC statistics. Within these 1118 pairs, our method identified 155 pairs while LDSC identified 49, at the same p-value significance of 1.0e-3 after Bonferroni correction (for our method, we required that the p values from both directions pass the significance threshold). The two predictions have 28 pairs overlapping (p-value for the overlap is 1.0e-12) (Dataset S4). This suggests that although gene-based method has higher sensitivity, the two methods are complementary. We found that HDL, an extension of LDSC with improved sensitivity[11], can detect the relation between CD and RA with marginal significance (p-value 4.5e-3), while our method detect this pair with much stronger significance (p-value <=5.6e-11).

One limitation of the gene-based method is that it does not predict the direction by which the two traits are correlated, while LDSC does predict a direction. Predicting direction by gene-based approach is complicated by several factors: 1) the p-value of association between a gene and a phenotype is typically supported by multiple SNPs, and when considering the relative sign between the effect on gene expression and the effect on phenotype, these SNPs can have either concordant or discordant directions. This is a previously observed phenomenon with an unknown mechanism[19] and makes it difficult to assign a direction for the gene-phenotype association, although it can be analyzed case by case. 2) the directional information for SNPs are not always available. We do not have directional information for most of the eQTL data we used except for GTEx. GTEx is systematic, but due to sample size limitation does not give reliable trans-eQTL information. The simplicity of the gene-based method is that it only needs p-values of association for the eQTL and GWAS data. Future work is needed to factor in directional information as more systematic and larger scale eQTL data become available.

A key step in our gene-based method is to translate the SNP-phenotype association profile to the gene-phenotype association profile. For this purpose, we developed the Sherlock-II algorithm, which is a second generation of the Sherlock algorithm we developed before. Sherlock-II kept the basic spirit of Sherlock in that it integrates information from all the SNPs (SNPs with both strong and weak GWAS p-values, and both cis and trans to a gene), but is more robust and efficient than Sherlock. The integration over all SNPs is essential for translating the global SNP-phenotype profile to gene-phenotype profile while preserving as much information as possible. This is important for detecting signals in the gene-phenotype association and for the success of the gene-based similarity measure.

A number of elegant GWAS vs. eQTL alignment tools have been developed, most of them focus on the colocalization between strong GWAS peaks and cis-eQTL peaks. For example, eCAVIER uses a Bayesian model to colocalize eQTL and GWAS peaks by considering local LD structure[18]. SMR computes the effect size from gene to phenotype and tests for consistency across strong cis-eSNPs that influence the same gene[19]. Both methods start with a strong GWAS/eQTL peak passing genome-wide significance threshold and analyze the region around it, leading to high specificity of the predictions. However, they infer genes based on mostly a single strong signal instead of integrating signals from multiple SNPs. Compared with these two methods, Sherlock-II detects gene-phenotype association with higher sensitivity, especially when the signals in the original GWAS are weak. For example, from the autism GWAS data, Sherlock-II identified 10 genes with FDR<0.1 (Figure S2), while eCAVIAR and SMR could not identify any gene because none of the autism GWAS signals passed the genome-wide significance threshold[35]. We found literature support for several of the 10 genes, including ENPP4, LEPR, and MTHFR [35, 36, 38, 63].

It is important to note that the data we obtained are co-associations between genes and phenotypes (supported by multiple GWAS and eQTL SNPs), not necessarily causal relations. When a gene G is found to be co-associated with phenotypes A and B, there are several different possible scenarios, including 1) G influences both A and B, and 2) G influence one of the phenotypes which in turn influences another phenotype. In some of the cases we discussed, such as the co-association of genes involved in Cancer and AD, it is plausible that 1) is the correct model, while for other cases such as CKD vs. cholesterol connection, model 2) is more likely (G influence CKD which in turn influence cholesterol). In many cases it might be difficult to distinguish the models, due to the lack of mechanistic understanding on how the genes connect to either of phenotypes. Clearly more experiments/analyses are required to sort out the causality for the genes/pathways we identified to be co-associated with many trait pairs.

## Methods

### Measuring genetic overlap between two traits using their gene-phenotype association profiles

#### Vector representation of the gene-phenotype association profile

We first translate the SNP-phenotype association profile (p-values of association for all the SNPs with the phenotype) into a gene-phenotype association profile (p-values of association for all the genes with the phenotype) using the Sherlock-II algorithm (see the last section in Methods). To measure genetic similarity between two phenotypes, we represent the association profile of a phenotype *t* by a vector of dimension *N* (the total number of genes in the genome), *V*(*t*) = [*Z_1_*(*t*), *Z_2_*(*t*), …, *Z_N_*(*t*)] where *Z_i_*(*t*) can be calculated from the p-value of association between gene *i* and trait *t*, by inverting the following equation: 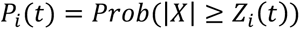, where *X* is a random variable with standard normal distribution. This p-value to z transformation is necessary to emphasize the contributions of genes with small p-values (indicating significant association between the expression level of the gene and the phenotype) when measuring the distance between a pair of gene-phenotype association vectors. The z score is unsigned, since we have p-value of association for the gene calculated from Sherlock-II without direction (see Discussion). We experimented with other types of vector representation, e.g., −*log*_10_(*P_i_*(*t*), and find very similar results (Figure S5)

#### Gene-based genetic overlap score between two phenotypes

Using the vector representation for the traits, we define a genetic overlap score between a pair of traits t1 and t2 as 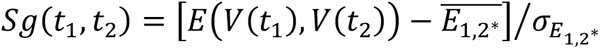 is the Euclidean distance between the two gene association vectors *V*(*t*_1_) and *V*(*t*_2_), 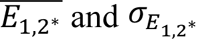 are the mean and standard deviation of a background distribution of Euclidean distances, generated by randomly permuting the components of the second vector (denoted as 2^∗^). A pair of phenotypes with co-associated genes are expected to have similar association vectors, and will give rise to smaller distance and more negative overlap score Sg. The normalization by the background distribution (measured as the number of standard deviations away from the mean) is essential as it takes out the contribution to the distance due to the features of each individual vectors, since we are interested in the overlap between the two vectors, not the vectors themselves. The background distribution is generated by 10^4 randomly permuted *V*(*t*_2_). This number of permutation is sufficient as the calculated mean and standard deviation converge to a value that no longer change with more simulations. We experimented with different distance metric (such as Manhattan distance) and found very similar results (Figure S6).

#### Assessing the statistical significance of the genetic overlap score

To assess the statistical significance of an observed genetic overlap score Sg between a query GWAS_1 and a hit GWAS_2, we generate an ensemble of random GWASs (GWAS_2R) with the same power as the query GWAS (see below), and use this ensemble to perform parallel calculations to generate a background distribution of overlap scores Sg(V1, V2R) between the query GWAS and the random GWASs (Figure 1). We then use this background distribution to normalize the score Sg(V1,V2), leading to a z score 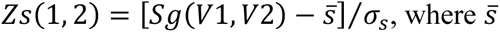 and σ*_s_* are the mean and standard deviation of the background distribution. This second normalization is necessary as the score Sg(V1, V2) does not follow z distribution even when one of the GWASs is a random GWAS (with signal randomly distributed to different SNPs), and the distribution further shifts away from z distribution as the power of the random GWAS increases (Figure S4). This is because the components of the two vectors are correlated even between a real GWAS and a random GWAS, due to the structure of the eQTL data. E.g., pleiotropic eSNPs will lead to the correlated fluctuation of the p values of their target genes, and such correlation is stronger for higher powered GWAS. By performing this pair specific second normalization, we take into account the dependence of the Sg on the power of GWASs being compared, resulting in a background distribution that follows z distribution (Figure S4). We then calculate a p value based on Zs(1,2) using z distribution.

To generate the ensemble of random GWASs (GWAS_2R), we first simulated a large number of random GWASs with different power (which we quantify as the percent of SNPs passing 10^(-6) pvalue cutoff). We simulated realistic random GWASs using imputed genotypes (version 3) from UK Biobank (UKBB) genotype data[64, 65]. UK Biobank is a large-scale biomedical database and research resource containing genetic, lifestyle and health information from half a million UK participants.

We took the following steps to prepare a set of 276,122 genotypes at 491,639 SNPs. We implemented these data preparation steps using PLINK version 2.00a3.1LM 64-bit Intel (19 May 2022) [66, 67]. To limit confounding due to population structure, we considered only genotype samples marked as “White British”[68] (based on a principal components analysis of the genotypes[64]). We then removed any samples from the “White British” subset that met one or more of the following criteria: mismatch between self-reported and genetic sex; outlier based on heterozygosity and/or rate of missing genotypes; and having at least one close relative in the same data set (based on the UK Biobank’s kinship and “relatedness” calculations). We selected SNPs if they were included in the list of 503,311 SNPs from the 69 GWASs that aligned to our eSNPs in the combined eQTL data. We additionally filtered out any SNPs that were not bi-allelic, or had minor allele frequencies (MAFs) less than 1% (specifically, we called PLINK with the following options: --snps-only --max-alleles 2 --rm-dup exclude-all).

We used these genotypes to simulate phenotypes with different number of causal SNPs (NS) and different population sizes (PS). We chose NS to be 100, 500, 1000, 3289, and PS from 10% to 100% of the UKBB, in 10% increment. This yielded a total of 40 combinations of NS and PS. For each of the 40 combinations, we simulated 100 random GWASs by picking NS random SNPs with their effect sizes sampled from the distribution of the effect sizes from the real GWAS data of a highly polygenic complex trait [69]. For a given set of causal SNPs and their associated effect sizes, we generated a continuous phenotype value for each individual given its genotype, using a linear additive model with a Gaussian noise term. Then the GWAS summary statistics was calculated from the resulting genotype – phenotype pairs using the standard procedure for analyzing quantitative traits[70]. We chose the standard deviation of Gaussian noise to be 1.8, so the percent of variance explained by the causal SNPs (which is ∼25%) as well as the power of the GWAS match those of the height GWAS when we use NS=3289 and PS= 100%. In total, we simulated 4000 random GWASs using the UKBB genotype data, the highest power of which matches that of the height GWASs, with the range of the power covering those of the 69 real GWAS data we used in this study.

Using the 4000 simulated random GWASs with different power, we then generated the random GWAS ensemble (GWAS_2R) matching a hit GWAS, by taking the top 200 random GWASs that best match the power of the hit GWAS (based on the percent of SNPs passing 10^(-6) pvalue threshold). Typically, the power of the matched random GWASs is within +/-15% of that of the hit GWAS. This ensemble was then used to normalize the score Sg(V1, V2) to derive the final zscore of the genetic overlap Zs(1,2).

To test this normalization scheme, we used 69 real GWASs as query against 200 random GWASs (randomly selected from the 4000) and found that after the second normalization, the distribution of score Zs followed the standard normal distribution for both high-powered and low-powered random GWASs (Fig S4B).

### Partial-Pearson-Correlation Analysis (PCCA) to identify sets of functionally related genes that contribute to the genetic similarity

We employed a simple statistical approach to detect gene ontology terms that contribute significantly to the genetic similarity between two different phenotypes. Gene Ontology gene annotations were downloaded from Ensembl. We removed the GO terms with less than 5 genes (not enough statistics) and more than 100 genes (too non-specific) from the analysis, after which 6796 GO terms were used. For a specific pair of phenotypes, we calculate a Partial-Pearson-Correlation-Analysis zscore (*Z*) for each of the 6796 GO terms using the following formula (previously developed for analyzing similarity between two gene expression profiles [15])

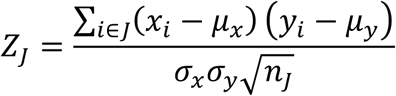

where *x_i_* and *y_i_* are the z values associated with the two phenotypes for gene *i* (calculated from the SherlockII p-values for gene-phenotype association), *μ_x_*,*σ_x_* and *μ_y_*,*σ_y_* are the mean and standard deviation for x and y respectively (calculated from all genes), J is the gene set defined by the GO term, and *n_j_* is the number of genes in the GO term. This formula is similar to the one for calculating Pearson correlation (with a normalization), except that the summation is partial – only for the genes in the subset. We argued that the null distribution for should be standard normal distribution, and verified that this is indeed the case by randomly sample all phenotypes pairs and generate for all the GO terms (Figure S8).

### Hierarchical clustering of phenotypes

To performed hierarchical clustering of traits based on their pairwise similarity scores, we first map the similarity score to a distance between 0 and 1 using the following equation: 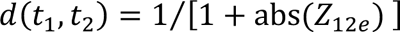, where Z_12e_ is the average of *Z*_12_ and *Z*_21_. For pairs with large negative z scores (with strong genetic overlap), the distance approaches 0. We then apply hierarchical clustering algorithm to this distance matrix, using average distance to measure the distance between composite nodes. We verified that the structure of the tree is not sensitive to the way the distance between composite nodes is measured. E.g, using weighted distance yielded a very similar tree (Figure S9)

### Sherlock-II algorithm: translating SNP-phenotype association profile to gene-phenotype association profile

Sherlock-II detects the association between an intermediate molecular trait (IMT) and a phenotype by comparing the GWAS SNPs with the QTL SNPs of the IMT (eQTL or metabolite-QTL), with the assumption that if an IMT is causal to the phenotype, SNPs that influence the IMT should also influence the phenotype, thus the QTL profile of the IMT should have significant overlap with the GWAS profile of the phenotype. To test the significance of the overlap, Sherlock-II first selects the QTL SNPs of an IMT that pass a threshold, aligns these SNPs to the corresponding SNPs in the GWAS, tags QTL SNPs in independent LD blocks, and then test whether the GWAS SNPs aligned to tagged QTL SNPs have p-values significantly better than those picked at random (see Figure S10 for an overall flow diagram of Sherlock-II).

When testing the significance of a selected set of loci in a given GWAS, Sherlock-II uses a simple scheme to compute a score for the n p-values of the GWAS SNPs that aliged to tagged eSNPs. Starting from an empirical background distribution for n=1, generated from the p values of GWAS SNPs aligned to tagged eSNP for all the genes, Sherlock-II uses a convolution-based approach to construct an empirical null distribution of scores involving n p-values drawn randomly from the background distribution for n=1. This test will be performed repeatedly for all intermediate molecular traits (e.g., all the genes from eQTL data) in a study. If dependence between the sets of IMTs is detected — for example, due to pleiotropic loci that regulate many genes — a provision of adjusting the null distribution is included. Through extensive testing and simulation, we show that our empirical strategy produces p-values that are highly resistant to inflation (Figure S12). In this section, we describe the individual components of the Sherlock-II method in detail in the order in which they occur.

#### Identifying Tag SNPs for Independent Blocks

Sherlock-II statistics is based on the assumption that the collection of SNPs tested are tagging independently segregating haplotype blocks in both the GWAS and eQTL cohorts. Understanding the true block structure of the cohorts is critical to accurately estimating gene significance. The inadvertent inclusion of dependent blocks can inflate the significance of genes by over-counting true associations; conversely, a conservative approach that enforces wide separation between tagging SNPs may miss many independent loci. The preferred approach to identifying independent blocks requires genotypes for both the GWAS and functional data to define block boundaries based on regions of rapid decay in linkage-disequilibrium (LD). However, since the difficulty of obtaining genotypes across a large number of GWAS studies makes this impractical, we constructed a database of LD using Caucasian cohorts from the 1000

Genomes Project. We use PLINK (v1.07) to compute r^2^ linkage between common SNPs for 379 individuals in the CEPH, CEU, TSI, GBR, FIN, and IBS cohorts of 1000 Genomes release v3.20101123. This permits the alignment of associated loci in the functional data to corresponding SNPs in GWAS data, and it enables the identification of independently-segregating tag SNPs in the combined data for use in the statistical test.

We first match the GWAS SNPs to all the loci present in a given functional data set that pass a specific threshold. For the eQTL data used here, the nominal threshold is typically eSNP p-values < 10 ^-5^. If matching SNPs are not present in both data sets, we use the closest GWAS SNPs in r^2^ LD > 0.85, if available. We then use an iterative approach to identify tagging eSNPs in independent LD blocks. For a given gene, after the alignment step, we start by tagging the most significant eSNP (with the best p value), removing all the other eSNPs in LD with this eSNP (defined by r^2^>=0.2), and then goes to the next most significant eSNP, and repeat the procedure until all the eSNPs are either tagged or removed. At the end of this procedure, any pair of tagged eSNPs will have r^2^< 0.2. We then use the GWAS p values of the SNPs aligned to the tagged eSNP for the subsequent statistical test. This same test is done for all the genes. Since discrepancies between the various cohorts — GWAS, functional, and linkage — are inevitable, a minimum 100 kb distance between tag SNPs is enforced. In addition, we exclude SNPs from the human leukocyte antigen (HLA) region between 6p22.1 to 6p21.3 due to its complex linkage structure.

We checked that our results are not sensitive to the choice of the block LD cutoff *r*^2^ (default 0.2) and the alignment cutoff LD r^2^ (default 0.85). The results do depend on the eQTL cutoff threshold, with 10^(-5) (the default value) generally giving better signals (more significant Zs) than the more stringent cutoff 10^(-6), suggesting that there are informative eSNPs with pvalue > 10^(-6) (Figure S11 and Data S5). In general, a highly significant pair remains highly significant with different parameter choices (Figure S11).

#### Core Statistical Method

Once a set of SNPs related to a specific molecular trait is identified, we compute a score s by simply combining their GWAS log p-values:

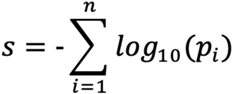

This converts our collection of SNPs into a scalar quantity that, when referenced against the appropriate null distribution, indicates the statistical significance of selecting this subset from the pool of all independent GWAS loci. Our approach is analogous to Fisher’s combined probability test using the scoring function 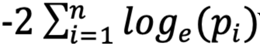 which follows a *X*^2^ distribution with 2n degrees of freedom when p-values are drawn from a uniform distribution on (0,1). With our method, instead of assuming a particular form for the distribution of GWAS p-values, we use discrete convolution to compute an empirically-derived distribution of scores when combining n p-values. Since s represents a specific score, we let S represent a random variable of all possible scores. We form the probability mass function (PMF) of S using bins of width b (log scale), where the probability of scores in the range 0≤ S < b is the first element, b ≤ S < 2b is the second element, and so forth. Thus, 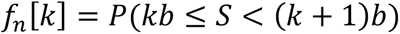 is k-th element of the PMF for a score S comprising n independent GWAS p-values. For the simplest case of scores involving only a single SNP, the PMF <u>*f*_1_ is essentially a normalized histogram of p-values for all independent (unlinked) SNPs in the GWAS that aligned to a tagged eQTL SNP across all genes</u>:

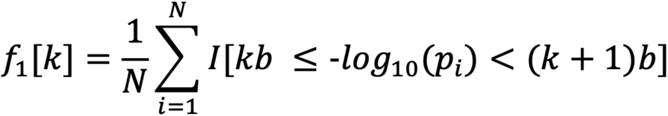

where I is the indicator function, N is the total number of SNPs, b is the bin width, and k is the array index within the histogram. For computational efficiency, the minimum GWAS p-value is typically truncated at 10 ^-9^, well below genome-wide significance, yielding 900 bins when spacing b = 0.01 is used. This single-locus PMF forms the basis from which null distributions for any arbitrary number of functionally-related loci are formed. Since the sum of two independent random variables has a PDF equal to the convolution of their individual probability distributions, scores involving two GWAS p-values have PMF 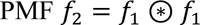. In the general case for any value of n > 1, the PMF *f_n_* is formed from n-1 recursive convolutions of *f_1_*:

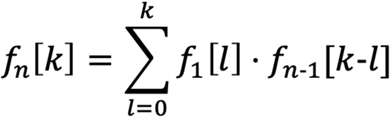

where again k is the array index of probability density function.

#### Computing the significance of a gene with score s

We used p-value to characterize the significance of the gene score s by summing the tail of the background distribution *f_n_*, where n is the number of independent GWAS loci supporting this gene. The tail is defined as the bins correspond to S that is great or equal to observe gene score s.

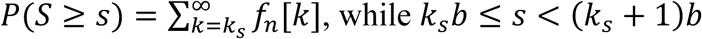

#### Correcting for Pleiotropic and Sampling Effect

Another source of inflation stems from a lack of true independence between molecular traits across a given functional dataset. For example, pleiotropic loci may appear to regulate hundreds of genes in the eQTL data for a given tissue. In our simulations, these loci may inflate the output test statistic due to chance alignment with significant GWAS SNPs; in our tests, this occurs at an appreciable level in perhaps five percent of the cases. Since our method permits the use of a unique distribution f_1_ for each p-value added to the score, it enables a simple scheme for reducing inflation by adjusting each distribution based on the number of molecular traits affected by the locus. For the eQTL example, this involves conditioning the GWAS distribution based on the number of genes influenced by a locus across the entire eQTL data set. When chance alignment of pleiotropic loci and significant GWAS SNPs occurs, the overrepresentation of small p-values is reflected in the null distribution for affected genes. Thus, in practice, the actual distribution incorporated into a Sherlock-II score with eQTL data is conditioned on the number of genes that are regulated by the same locus:

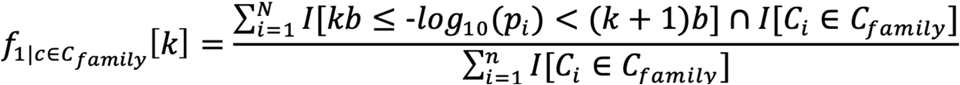

where I is the indicator function, N is the total number of SNPs, k is the index, b is the bin width, and the ith SNP is assigned gene count C_i_ while the locus in question has count c. *C_family_* is several non-overlapping sets, each including a range of gene count c. For example, *C_family6_* includes all SNPs that regulate 6 to 10 genes. Now the distribution contains only the p-values of SNPs that belong to the same count family. With this change, instead of computing the significance of a set of SNPs drawn from GWAS, our method computes the significance of a set of SNPs drawn given their individual pleiotropies count family. In our simulations, this results in a minor change in the significance and rank order of output genes in most cases but prevents inflation whenever chance alignment of significant GWAS SNPs and pleiotropic loci occurs (Figure S12). Specifically, we penalize SNPs regulating the expression of a large number of genes.

#### Reporting the statistics from Sherlock-II analyses

Unless otherwise explicitly stated, we reported the raw p value for gene/metabolite vs. phenotype association and we reported the number of gene-phenotype and metabolite-phenotype associations with Q<0.05.

## Data Availability

The genotype data of the UK Biobank can be accessed through procedures described on its webpage (https://www.ukbiobank.ac.uk/using-the-resource). Simulated data used in this paper can be obtained from the authors upon request. GWAS data analyzed in this paper are all published data and the relevant PMIDs were provided. All data produced by the Sherlock-II and the genetic overlap analyses will be made available upon publication.

## Code Availability

The source code for Sherlock-II and the genetic overlap analysis will be made available upon publication.

## Acknowledgement

The work was supported by NIH grants R21AG064357, R21AG071899, R01AG058742, and by a Chan Zuckerberg Biohub Investigator award and a BARI investigator award to HL.

## Author Contribution

JZ and HL conceived and designed the study. JG, CF, JZ performed computation. XH and PC performed simulations using UKBB genotype data. JG, JZ, HL analyzed and interpreted data. JG, JZ, and HL wrote the manuscript.

