## Supplementary figures and images for "Identifying the genetic basis and molecular mechanisms underlying phenotypic correlation between complex human traits using a gene-based approach"

### Figure S1

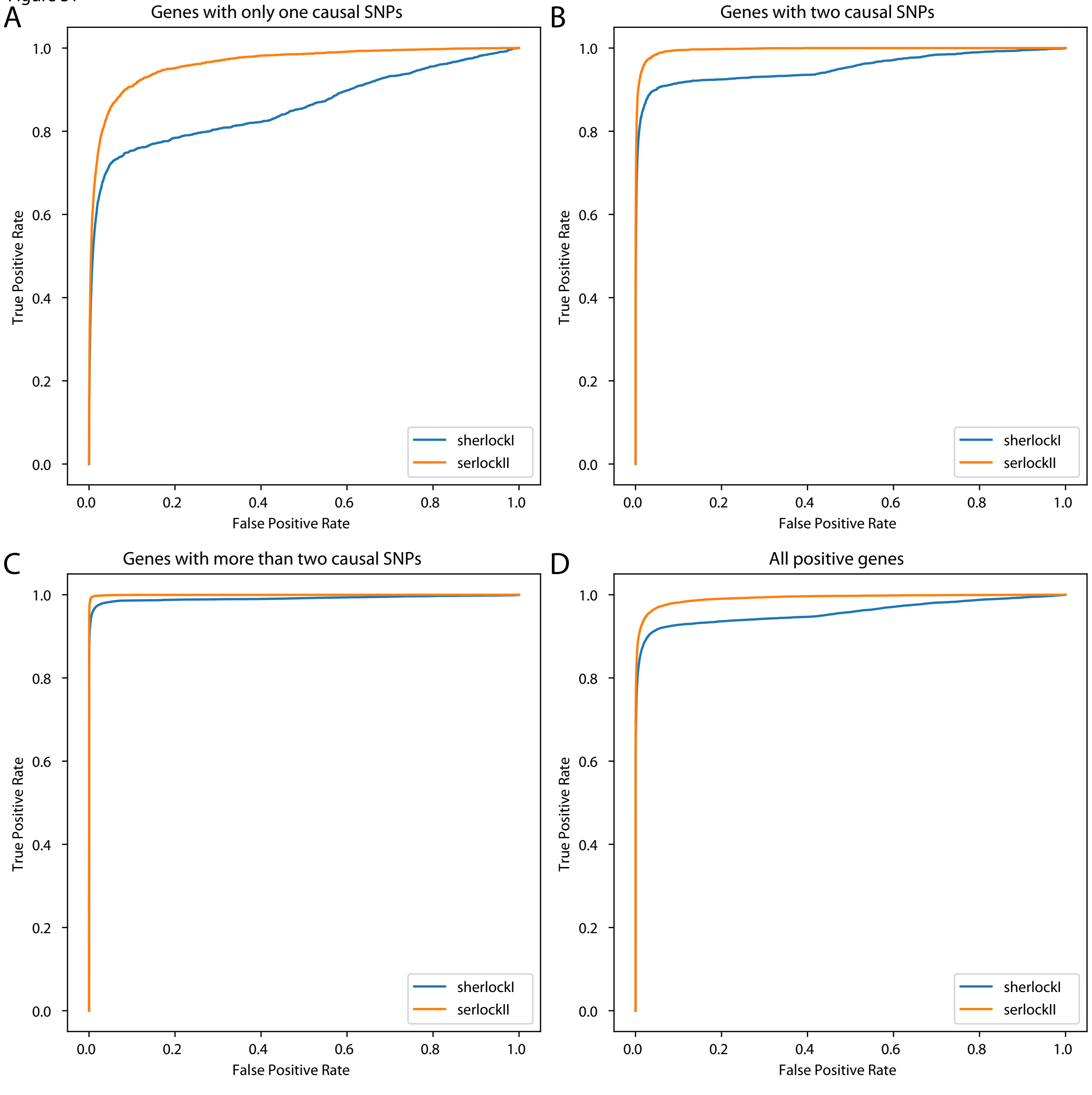

### Figure S2

Figure S2.

A

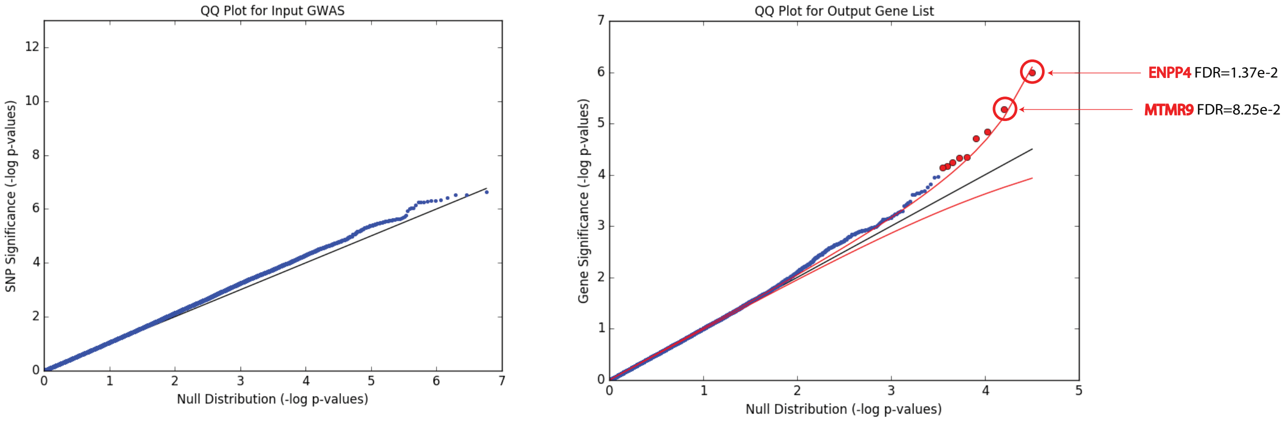


B

| Gene | BNIPL | ENPP4 | IL20RB | LEPR | MAPT | MTHFR | MTMR9 | RRP12 | SGK1 | UBE2Q2P1 |
|---|---|---|---|---|---|---|---|---|---|---|
| FDR | 4.76E-02 | 1.37E-02 | 1.22E-02 | 3.98E-02 | 8.05E-02 | 5.57E-02 | 8.25E-02 | 7.69E-02 | 7.94E-02 | 8.05E-02 |

### Figure S3

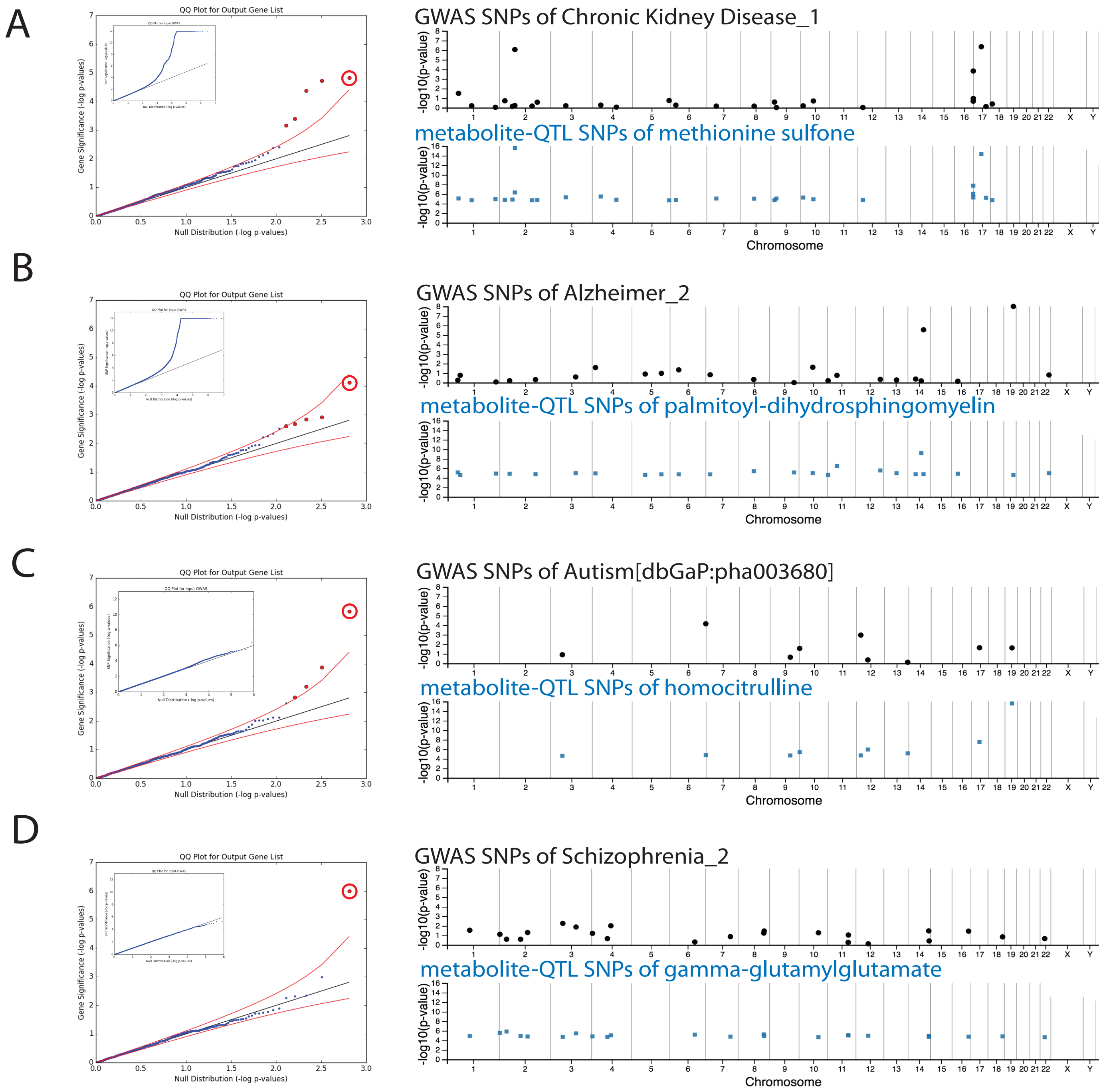

### Figure S4

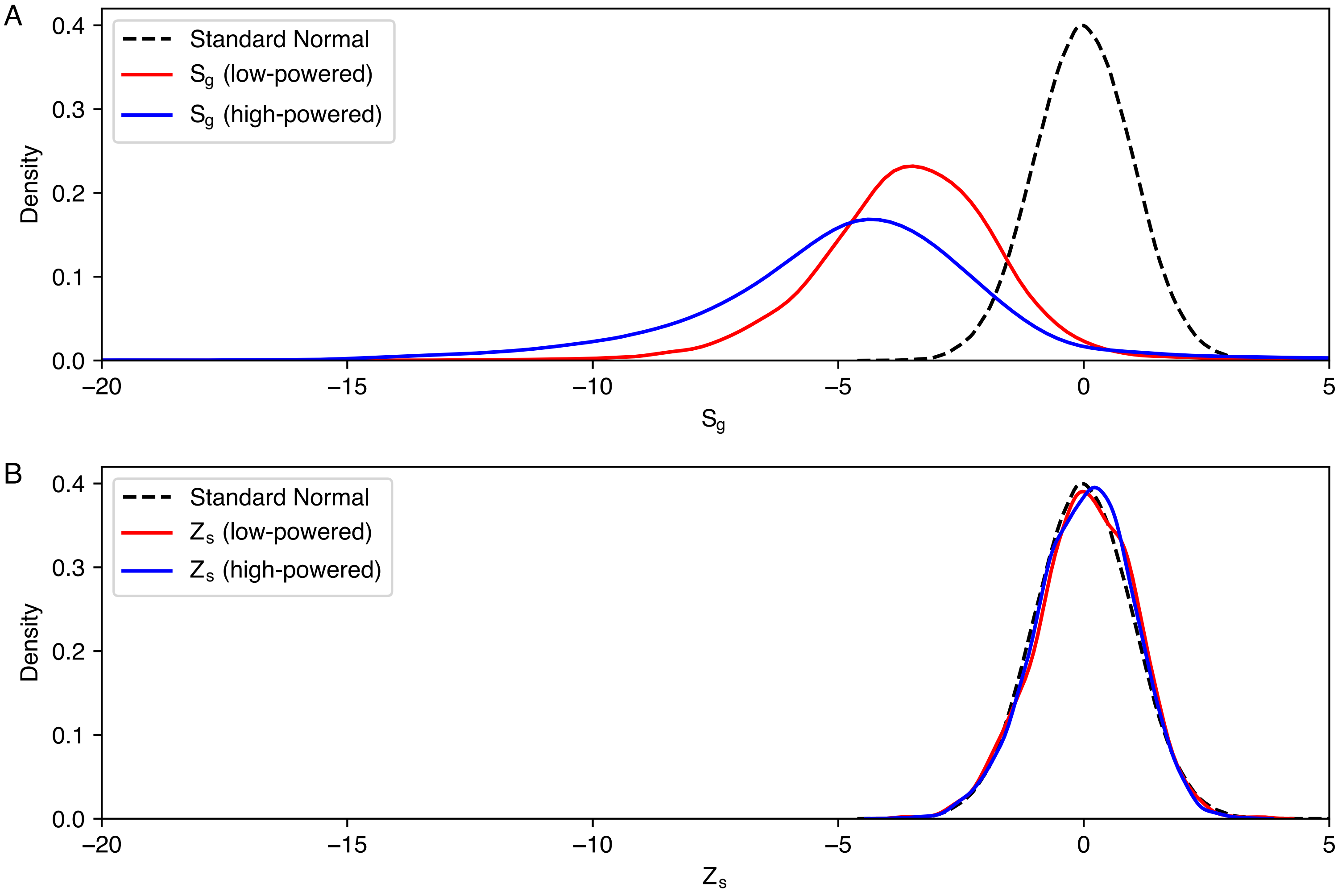

### Figure S5

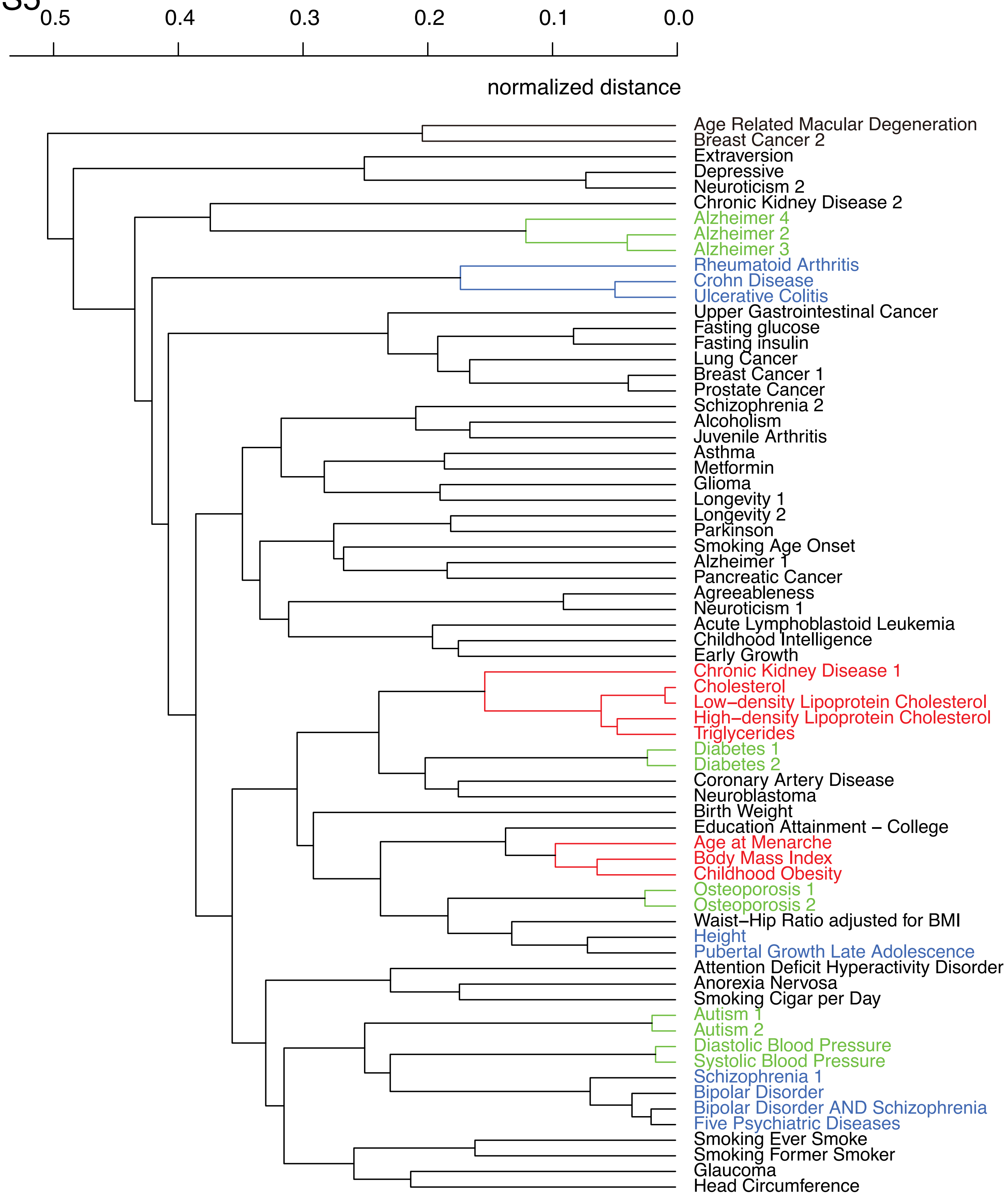

### Figure S6

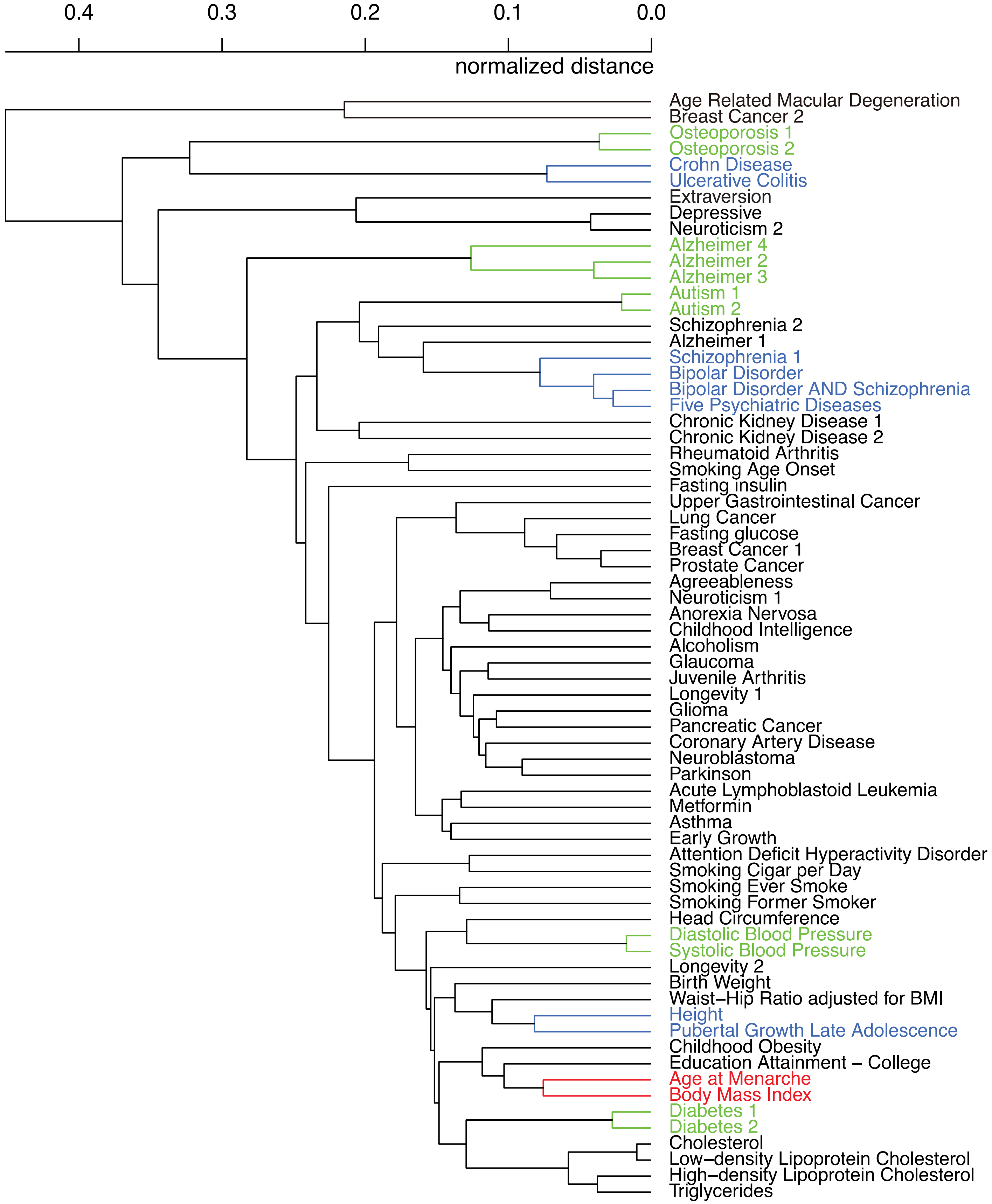

### Figure S7

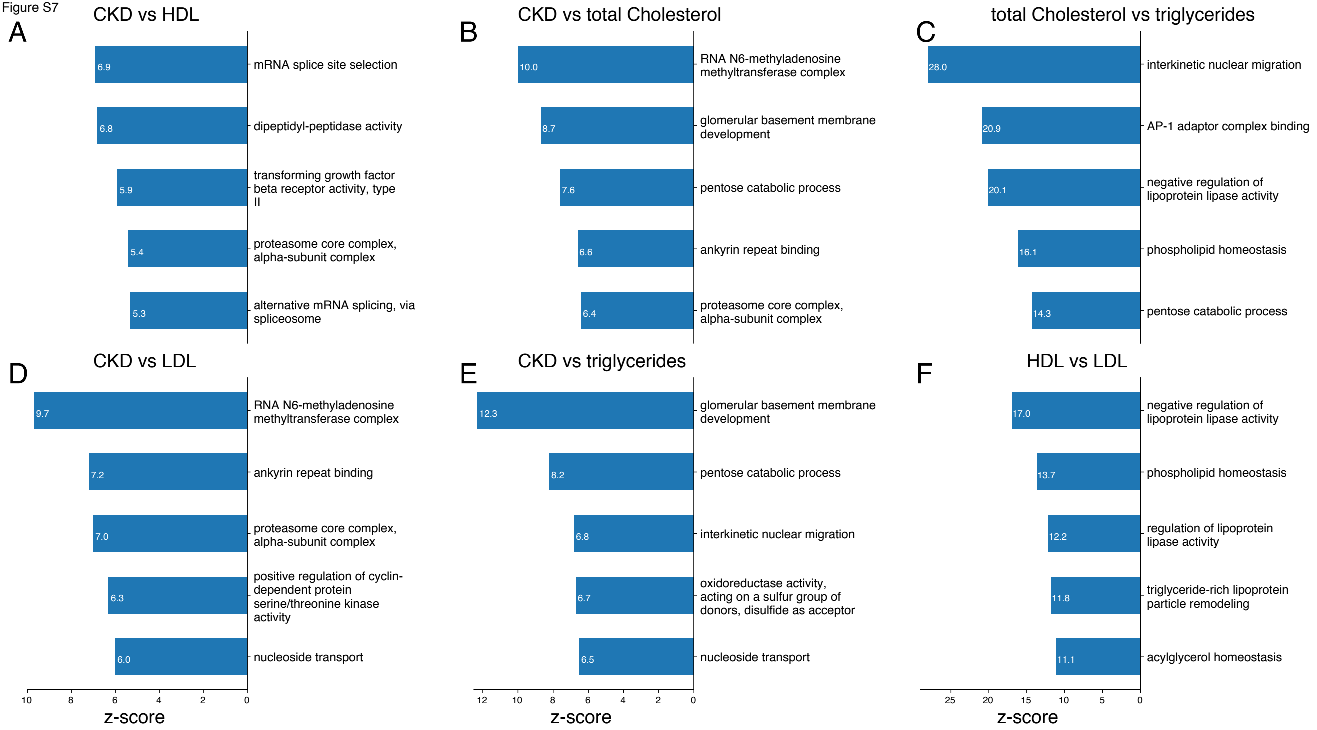

### Figure S8

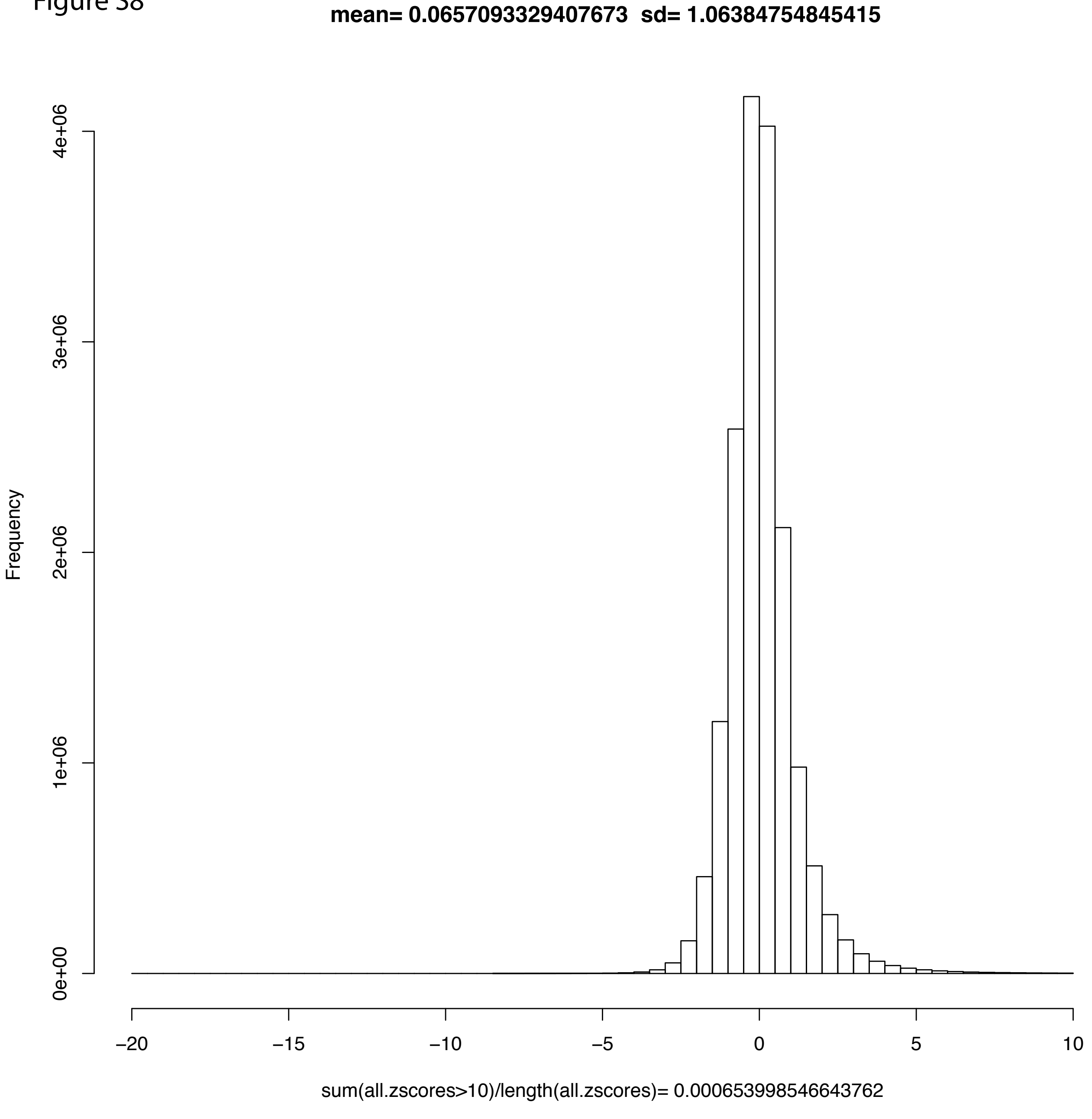

### Figure S9

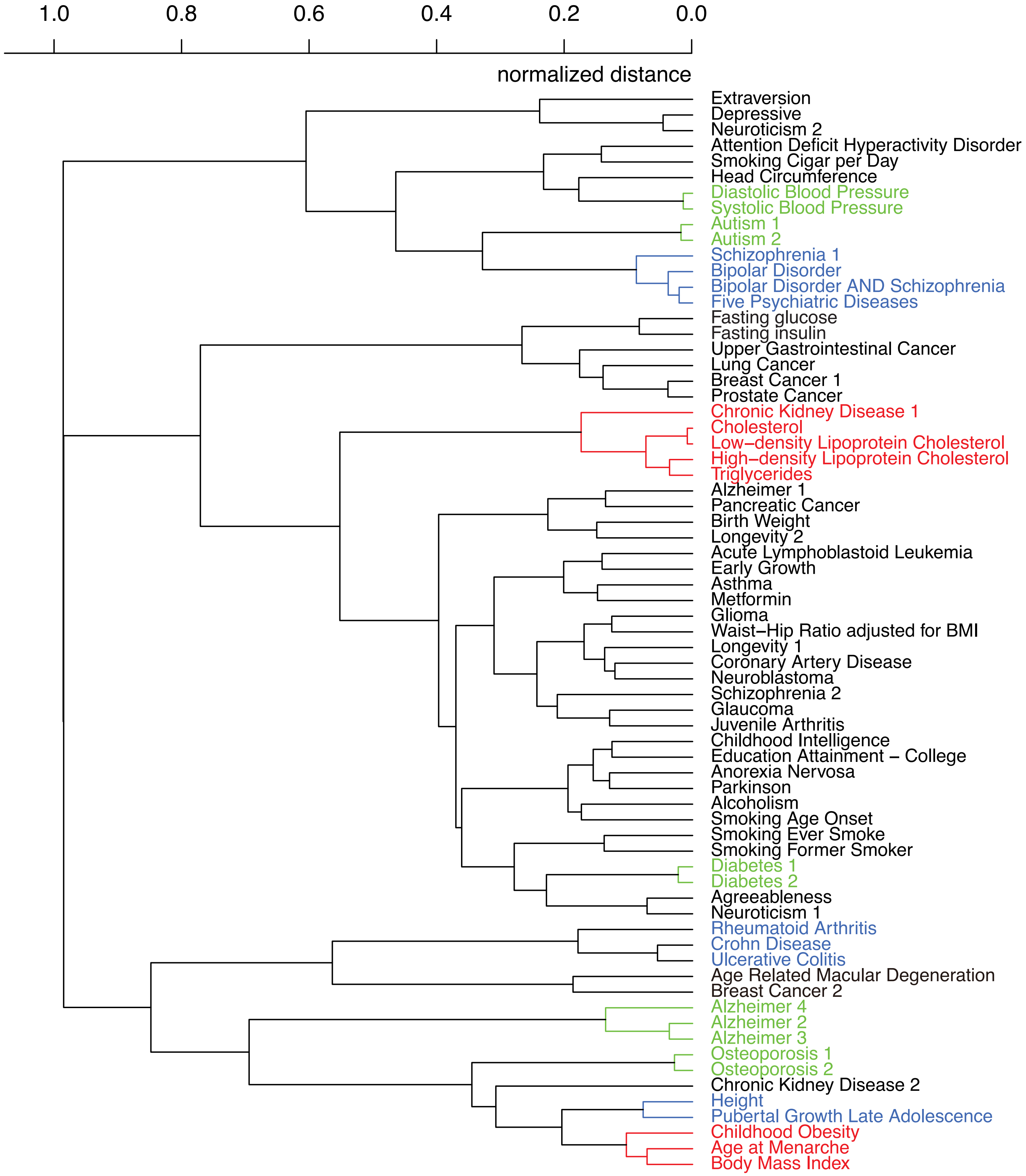

### Figure S10

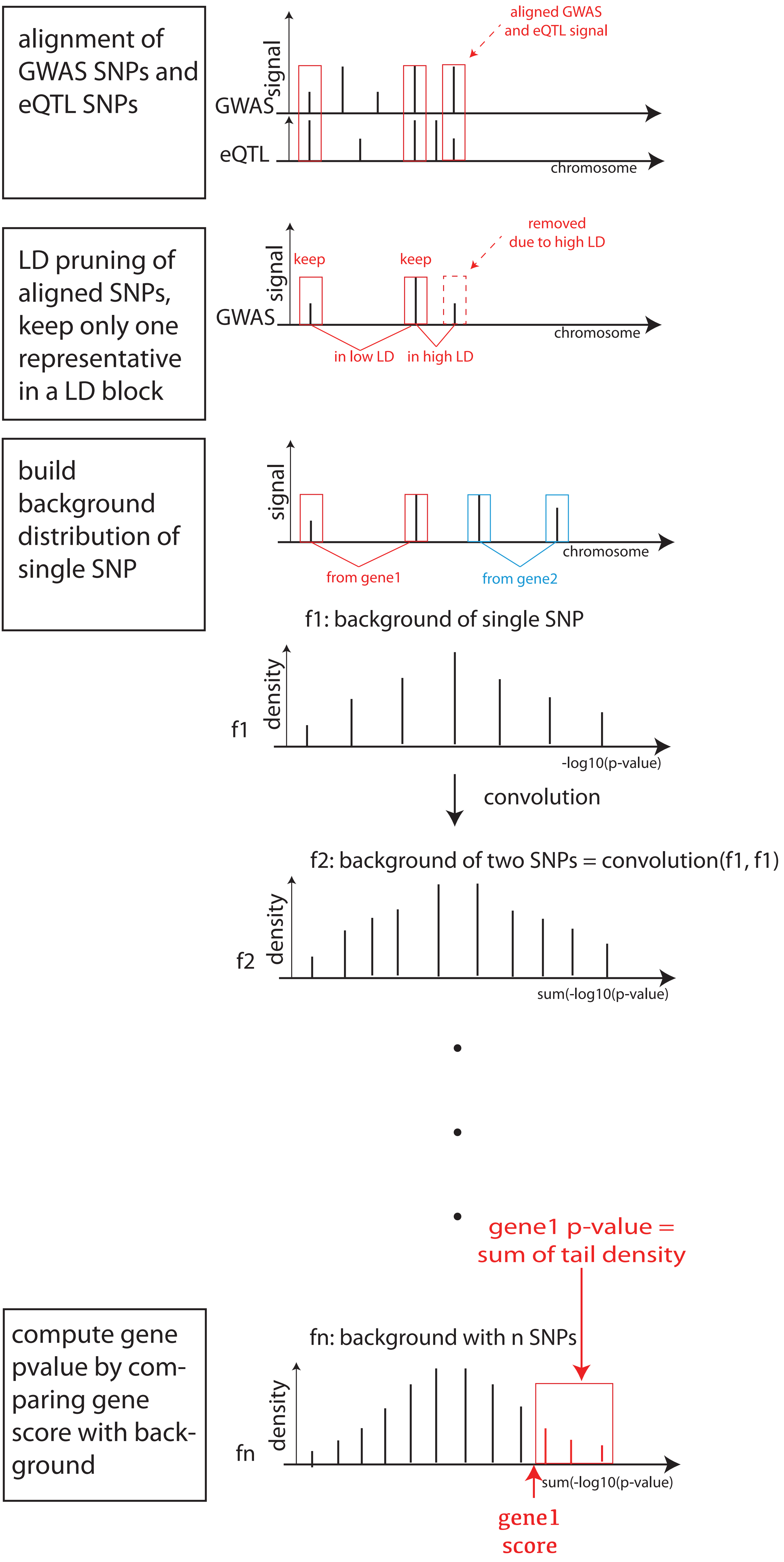

### Figure S11

Figure S11

#### Phenotype overlap score $Z_s$ from different SherlockII parameters

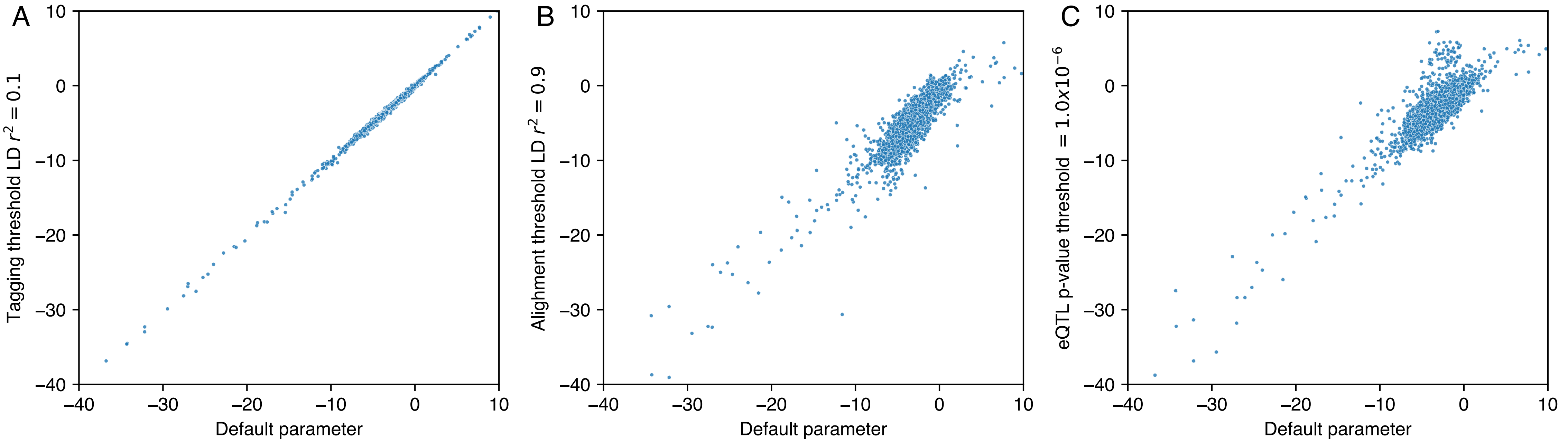

### Figure S12

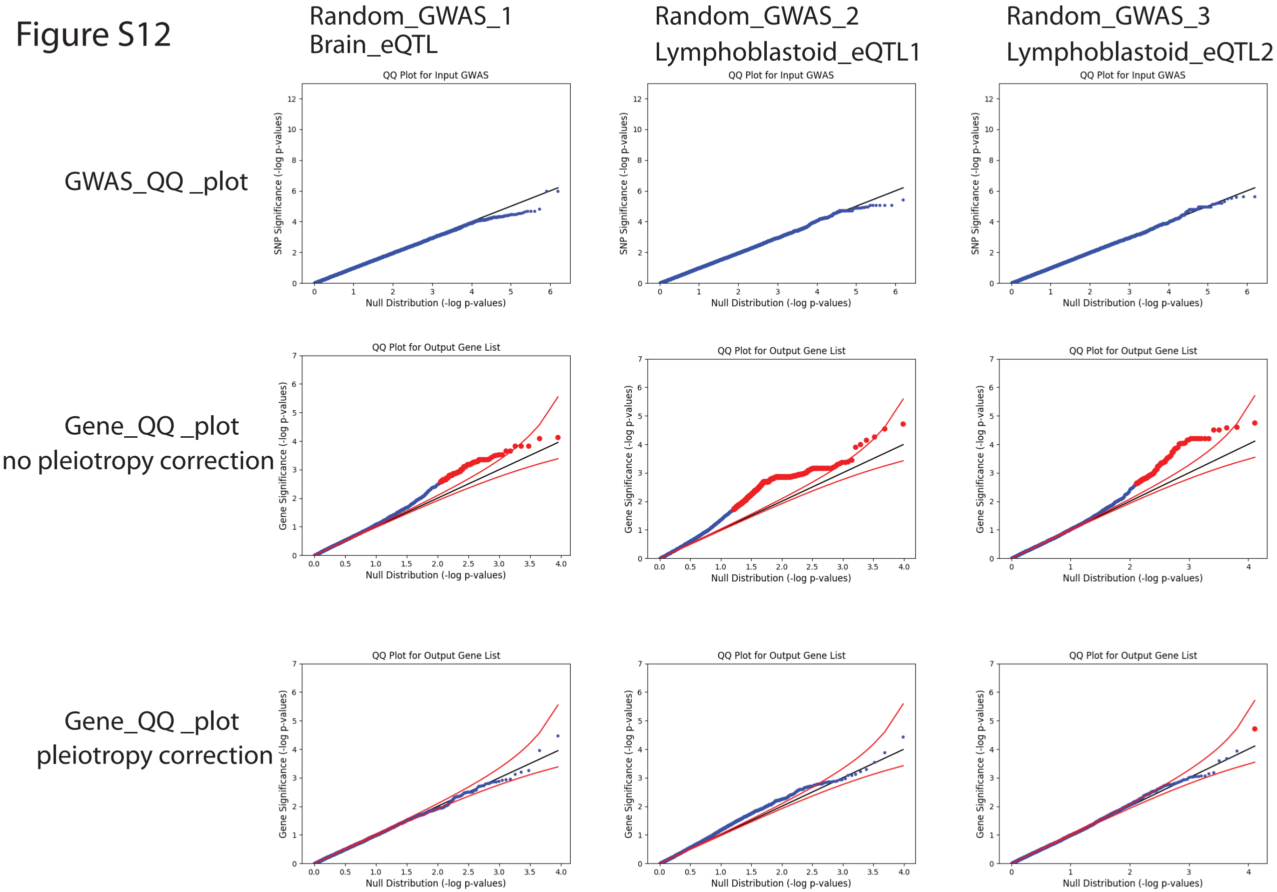
